# Microbial bioprospecting for benzoxazolinate-like molecules: unleashing the potential of genome mining

**DOI:** 10.64898/2026.08.17.745186

**Authors:** Sunaina Paliyal, Babanpreet Kaur, Latika Rao, Ardhendu Chakrabortty, Lovepreet Singh, Ishita Sehgal, Muskan Sharma, Dalwinder Singh, Vasvi Chaudhry, Shrikant S. Mantri

## Abstract

The benzoxazolinate moiety is a key functional group found in a few natural products (NPs), exhibiting diverse bioactivities, including antitumor, antibacterial, and cytotoxic activities. Despite their clinical importance, only a few bacterial strains and NPs have been reported harboring this rare bis-heterocyclic moiety, underscoring a largely unexplored chemical space. Here, we performed large-scale genome mining and identified 277 putative biosynthetic gene clusters (BGCs) across diverse bacterial hosts, including previously unreported bacterial genera and strains. The BGCs were grouped into three compound classes: benzoxazolinate, benzobactin, and ashimides based on sequence similarity network clustering. Bioactivity predictions of the identified BGCs revealed the predominance of antibacterial and cytotoxic potential, highlighting promising candidates for future experimental validation and functional studies. This study also presents a neural network-based bioprospecting model that efficiently detects rare BGCs encoding benzoxazolinate-containing molecules from genomic sequences. Overall, our findings expand the known repertoire of bacterial hosts with the potential to produce benzoxazolinate-containing NPs and provide a comprehensive framework for the discovery and identification of candidate BGCs.

**Importance:** This study helped uncover previously unknown bacterial hosts with the potential to encode benzoxazolinate-containing NPs through extensive genome mining. The findings suggest that benzoxazolinate-associated biosynthetic potential is more widespread than previously recognized and often overlooked by conventional annotation tools. We developed a neural network-based bioprospecting model to rapidly identify rare clusters in genomic and metagenomic datasets with high accuracy. Our work demonstrates a systematic strategy for uncovering cryptic gene clusters associated with benzoxazolinate-like metabolites across microbial genomes, thereby advancing a rational and scalable approach for future natural product discovery efforts.

## Introduction

Microbial bioprospecting has emerged as a promising strategy for exploring and identifying novel microbial species that encode bioactive secondary metabolites (SMs), which are widespread across diverse ecological niches and have industrial, pharmaceutical, and agricultural applications (Beattie et al., 2011; Perez Rojo et al., 2023). As antimicrobial resistance and cancer rates continue to rise, the search for novel, effective, and safer medications becomes essential and demands immediate attention. NPs are more chemically and biologically diverse than primary metabolites and serve as a rich source for the discovery of novel drugs to treat infectious diseases and cancer (Sekurova et al., 2019). Based on their biosynthetic pathways, these NPs are classified into major biosynthetic classes, including nonribosomal peptide (NRPS), polyketide synthase (PKS), terpene saccharides, and ribosomally synthesized and posttranslationally modified peptides (RiPPs) (Mantri et al., 2021). The genes encoding these biosynthesis pathways are physically clustered within a genomic region and together constitute biosynthetic gene clusters (BGCs) (Zdouc et al., 2025). These clusters are primarily composed of core biosynthetic genes that determine the biosynthetic class of SMs (Kim & Dettman, 2025), as well as additional, regulatory, and transport-related genes.

Recent advances in genomic and metagenomic data have enabled in silico bioprospecting methods to screen (meta)genomic datasets for target BGCs (Negri et al., 2022; Vuong et al., 2022; Ziemert et al., 2016). Computational tools such as BAGEL (de Jong et al., 2006), PRISM (Skinnider et al., 2017), and antiSMASH (Blin et al., 2023) provide a comprehensive pipeline for the detection and annotation of BGCs and are the gold standard in genome mining of BGCs; however, their reliance on predefined rules (e.g., antiSMASH) for BGC annotation limits their ability to detect novel and uncharacterized BGCs (Lee et al., 2020; Zhu et al., 2025). These genome mining approaches have led to the identification of a greater biosynthetic diversity across microbial genomes than the known repertoire of characterized BGC-SM pairs. These findings also indicate that the majority of chemical diversity remains unexplored in microbial genomes and necessitate the need for large-scale genome mining efforts to uncover the hidden microbial potential (Gavriilidou et al., 2022).

Among structurally diverse SMs, the enediyne family of antitumor antibiotics harbors the most cytotoxic NPs. One such promising candidate is C-1027 (Lidamycin, LDM), which features a rare benzoxazolinate moiety that contributes to its cytotoxicity against cancer cells (Cui et al., 2009; Xu et al., 1994; Zhen et al., 2009). The unique structural and biological significance of benzoxazolinate has drawn attention to NPs with this moiety as a valuable source of bioactive compounds. This moiety was first identified in *Streptomyces globisporus* C-1027, encoded by *sgc* cluster (Liu et al., 2002). Later, through genome mining, the *xsb* cluster was characterized in *Xenorhabdus szentirmaii* DSM 16338 (Y. Shi et al., 2022) (Fig. 1a). The *in vitro* functional characterization revealed the three-gene cassette responsible for the biosynthesis of benzoxazolinate in *X. szentirmaii,* comprising *xsbA* encoding FMN-reductase (dehydrogenase), *xsbC* encoding acyl-AMP ligase, and *xpzC* encoding 2-amino-2-deoxyisochorismic acid (ADIC) synthase. The ADIC synthase is derived from the phenazine biosynthesis pathway *(xpz*) and is located approximately 2 Mb away from the *xsb* gene cluster (Fig. 1a). Similarly, the *asm* gene cluster encoding cytotoxic ashimides A and B was characterized in *Streptomyces* sp. NA03103. In addition to the core genes involved in the biosynthesis of benzoxazolinate, the *asm* gene cluster has additional genes required for the biosynthesis of ashimides (Fig. 1b) (J. Shi et al., 2019). In addition to the discovery of benzoxazolinate, the identification of cytotoxic benzobactin, a derivative of benzoxazolinate, was highlighted through the identification of the *xvb* gene cluster from *X*. *vietnamensis* DSM 22392 (Fig. 1a) (Y.-M. Shi et al., 2022). ADIC derived from the shikimic acid pathway, serves as the branching point for the biosynthesis of benzoxazolinate and its derivatives. The chemical analysis of the cytotoxic benzobactin revealed the incorporation of two non-proteinogenic amino acid residues, 2-hydroxymethylserines, covalently linked to the benzoxazolinate moiety by *xvbB* encoding serine hydroxymethyltransferase (SHMT) (Y.-M. Shi et al., 2022) (Fig. 1c). At present, few known compounds contain the benzoxazolinate moiety as part of their core structure. Given the promising antitumor bioactivity associated with benzoxazolinate, extensive genomic datasets can be mined to identify analogous biosynthetic genes within the diverse genomes of microorganisms.

**Figure 1:**
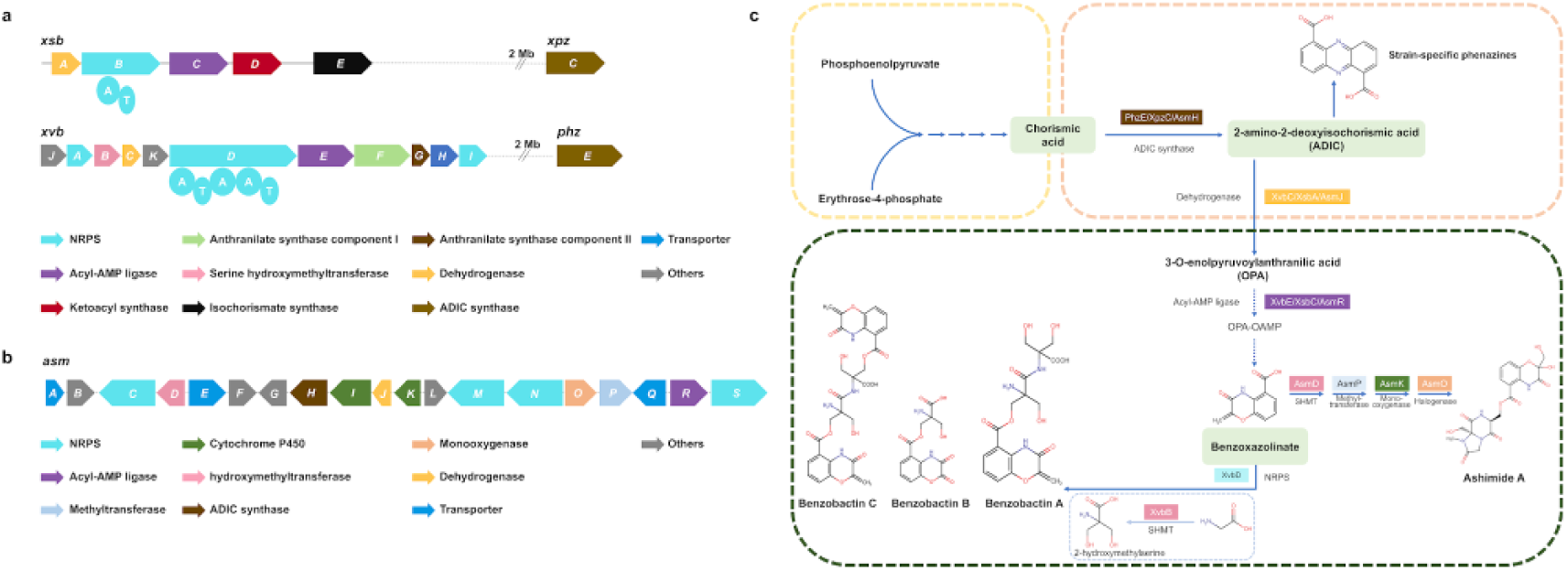
Biosynthetic genes involved in the biosynthesis of benzoxazolinate, benzobactin, and ashimides. a) *xsb* cluster from *X. szentirmai*i DSM 16338 encoding benzoxazolinate moiety and *xvb* cluster from *X. vietnamensis* DSM 22392 encoding benzobactin. b) *asm* cluster from *Streptomyces sp.* NA03103 encoding ashimides. c) Metabolic pathways illustrating branching points from chorismic acid, ADIC, and benzoxazolinate. ADIC serves as the key intermediate for the biosynthesis of phenazine and benzoxazolinate. Benzoxazolinate serves as a key building block for the biosynthesis of benzobactin and ashimide derivatives.

The structural diversification of these SMs is primarily catalyzed by tailoring enzymes, which modify the core scaffold generated by gatekeeper enzymes. The nascent metabolites undergo chemical modifications such as halogenation and methylation, generating bioactive and stable NPs. Currently, Minimum Information about a Tailoring Enzyme (MITE) database provides comprehensive information on SM-acting tailoring enzymes (Rutz et al., 2025). MITE is used as a reference database for enzyme annotation in the latest antiSMASH version (Blin et al., 2025). However, the database comprises only 202 active entries, underscoring the urgent need to identify and characterize novel tailoring enzymes within the bacterial BGCs in the context of the rapidly expanding discovery of BGCs, as these tailoring enzymes are critical for NP diversity.

Previous genome mining studies characterized the biosynthesis of benzoxazolinate-containing NPs in *Xenorhabdus*, *Pseudomonas,* and *Streptomyces* (J. Shi et al., 2019; Y. Shi et al., 2022; Y.-M. Shi et al., 2022). These studies characterized the core enzymatic machinery involved in the biosynthesis but revealed limited taxonomic distribution and biosynthetic diversity. Our study primarily focuses on large-scale, systematic genome mining to identify candidate BGCs encoding benzoxazolinate-containing NPs across diverse bacterial genomes (Fig. 2). We conducted a sequence-based database search and identified diverse bacterial hosts harboring putative BGCs. To determine their bioactive potential, we performed bioactivity predictions, which revealed diverse predicted bioactivities associated with identified clusters. To quantify patterns of genetic diversity, we analyzed gene cluster families (GCFs) for openness metrics, revealing that diversity is primarily due to gene reshuffling. To explore their ecological abundance, we conducted a pilot rhizosphere microbiome analysis that revealed distinct abundance patterns for identified bacterial strains. Finally, to enable systematic screening of rare biosynthetic clusters, we developed a neural network-based bioprospecting model that achieved high prediction accuracy.

**Figure 2:**
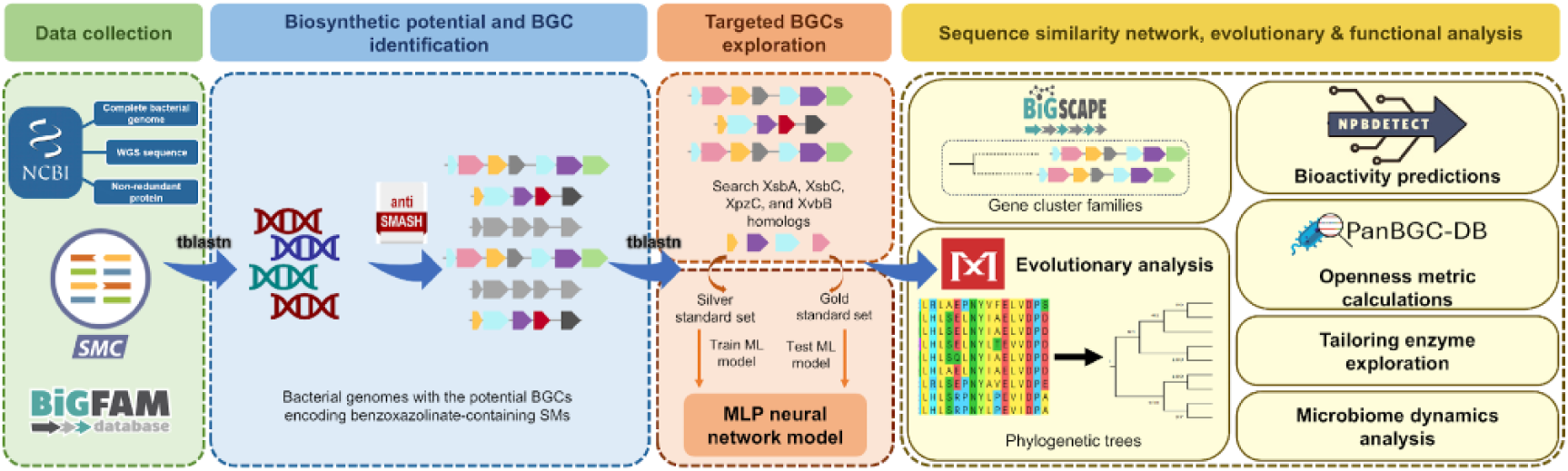
Integrated workflow for genome mining and BGC identification. An overview of the analytical pipeline employed for identifying potential benzoxazolinate-encoding BGCs. Data were retrieved from NCBI, the secondary metabolism collaboratory (SMC), and the BiG-FAM database. Candidate BGCs were identified using sequence similarity search, followed by developing a neural network model for bioprospecting benzoxazolinate-containing SM-BGCs. The identified BGCs were further analyzed for GCF classification, and subsequent evolutionary and functional analysis.

## Results and Discussion

### Identification of bacterial hosts with potential benzoxazolinate-like BGCs

To identify potential bacterial hosts encoding benzoxazolinate-containing SMs, publicly available bacterial genomes in the NCBI complete prokaryotic genome and whole-genome shotgun (WGS) database were mined. A total of 14,694 hits from the bacterial genomes dataset were screened, leading to the identification of a total of 302 bacterial genomes. Similarly, 629 hits were retrieved from the WGS dataset that showed significant similarity to the query proteins. Following overlap analysis, 37 WGS sequences were retained, comprising combinations of three (XsbA, XsbC, and XpzC) or all four (including XvbB) proteins (Fig. S1). Out of the 37, only 16 WGS sequences were successfully retrieved using the NCBI Batch Entrez utility. In the tBLASTn analysis of query proteins against the secondary metabolism collaboratory (SMC) database, 2,000 significant hits were identified. Overlap analysis identified 164 BGCs containing the combination of XsbA, XsbC, and XvbB and 241 BGCs containing XsbA and XsbC (Fig. S2). A comprehensive blastp analysis of the non-redundant (nr) protein database identified 28 bacterial hosts harboring homologs of target proteins.

### Identification of target BGCs from genomic datasets

The biosynthetic potential of the selected bacterial genomes was assessed using antiSMASH v7.1, which identified 4,759 BGCs spanning different biosynthetic classes. To identify the putative BGCs encoding benzoxazolinate-containing SMs and to remove the background noise, the resulting BGCs were used to generate a BLAST nucleotide database. In tBLASTn of proteins of interest (XsbA, XsbC, XpzC, and XvbB) against the generated nucleotide database, 659 significant hits were obtained. After the overlap analysis, 87 BGCs were retained, each containing three proteins (XsbA, XsbC, and XpzC) or all four proteins (including XvbB), involved in benzoxazolinate and benzobactin biosynthesis (Fig. S3).

antiSMASH annotations of the SMC database identified 404 candidate BGCs encoding benzoxazolinate-containing SMs, while the nr-protein database identified 21 potential BGCs encoding benzoxazolinate-containing SMs. Exploration of the BiG-FAM database identified GCF_13705 that contains 17 core members (dist ≤ 900.0) and six putative members (dist >900.0). These core BGCs are represented by four different genera: *Bacillus*, *Clostridium*, *Pseudomonas*, and *Xenorhabdus* (Table S1).

This comprehensive analysis identified 529 putative BGCs (Fig. S4, Table S2). After deduplication, 277 non-redundant unique BGCs were retained and subjected to further downstream analysis (Table S3).

### Distribution and diversity of benzoxazolinate-encoding hosts

Further exploration of the identified BGCs revealed 153 putative BGCs with the potential to encode benzoxazolinate-like molecules, 100 with the potential to encode benzobactin-like molecules, and 24 with the potential to encode ashimides-like SMs (Fig. 3). Out of the 277 BGCs, 135 (48.7%) were identified as previously unreported microbial species of known genera, 69 (24.9%) were from unreported genera, and 53 BGCs (19.1%) were from unreported strains of known species and the rest were known candidates (Fig. 3). Overall, the maximum abundance of unreported species, subspecies, and strains was within the *Streptomycetaceae* family (106), followed by the *Pseudomonadaceae* family (67) (Fig. 3). Metadata on available habitat and geographical distribution of genome mined bacterial strains were compiled (Table S3). These strains were reported from different countries, including China, USA, Japan, and India (Fig. S5), and were isolated from diverse environments, including terrestrial (soil), plant-associated, aquatic, and animal-associated habitats. A notable proportion of BGCs originated from unknown or unreported sources, highlighting the need to curate more detailed ecological metadata in genome databases (Fig. 3).

**Figure 3:**
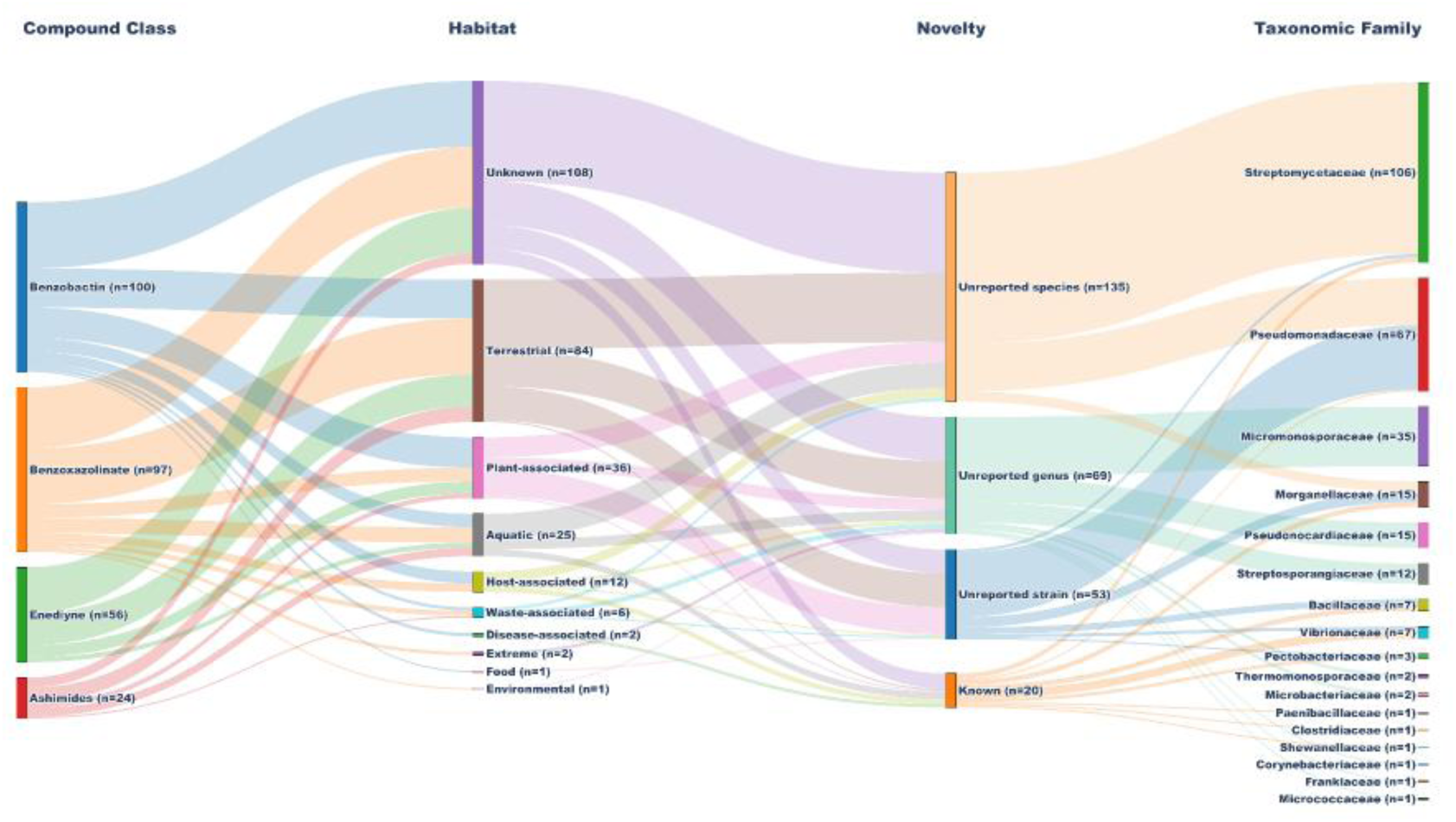
Distribution of 277 BGC encoding bacterial hosts across compound classes, ecological habitats, novelty status, and taxonomic families visualised as a Sankey diagram. The majority of bacterial hosts belong to unreported species and are predominantly associated with the family *Streptomycetaceae*. The width of each flow corresponds to the number of bacterial hosts associated with adjacent categories. Notably, 56 BGCs associated with benzoxazolinate biosynthesis harbored the conserved gene cassette involved in enediyne biosynthesis.

### Sequence similarity based organisation of candidate BGCs

The candidate BGCs identified across distinct bacterial genomes were analyzed based on sequence similarity using BiG-SCAPE v1.1.9 (Navarro-Muñoz et al., 2020). Out of 277 genome mined BGCs, 24 displayed cluster organization similar to ashimide-like BGC. These were grouped into nine GCFs, including five singletons. The presence of singletons indicates sequence divergence relative to reference and recovered ashimide-like clusters. One major GCF (FAM_02515) comprising 13 members clustered with MIBiG ashimide reference BGC, suggesting conservation of core biosynthetic architecture. Similarly, the 100 candidate benzobactin-like BGCs were organised into 21 distinct GCFs. Among these, eight GCFs were represented by singletons, and the remaining BGCs were distributed across multi-member families. As experimentally validated benzobactin clusters are absent from the MIBiG repository, direct comparison with reference clusters was not possible. The distribution of benzobactin BGCs across multiple GCFs highlights sequence diversity within the recovered benzobactin clusters.

Among 153 candidate benzoxazolinate-like BGCs, 56 BGCs were identified with conserved gene cassettes involved in the enediyne biosynthesis. Of these, 12 clustered with known MIBiG BGCs encoding the antitumor enediyne compound C-1027. The presence of known C-1027 clusters suggests a conserved core gene cassette involved in the nine-membered enediyne biosynthesis and highlights the potential to encode structurally related cytotoxic metabolites (Han & Seyedsayamdost, 2024; Yan et al., 2016).

The distribution of these BGCs in diverse biosynthetic classes (Fig. 4) highlights the current limitations of the rule-based classification system (discussed in Future outlook, recommendations, and limitations).

**Figure 4:**
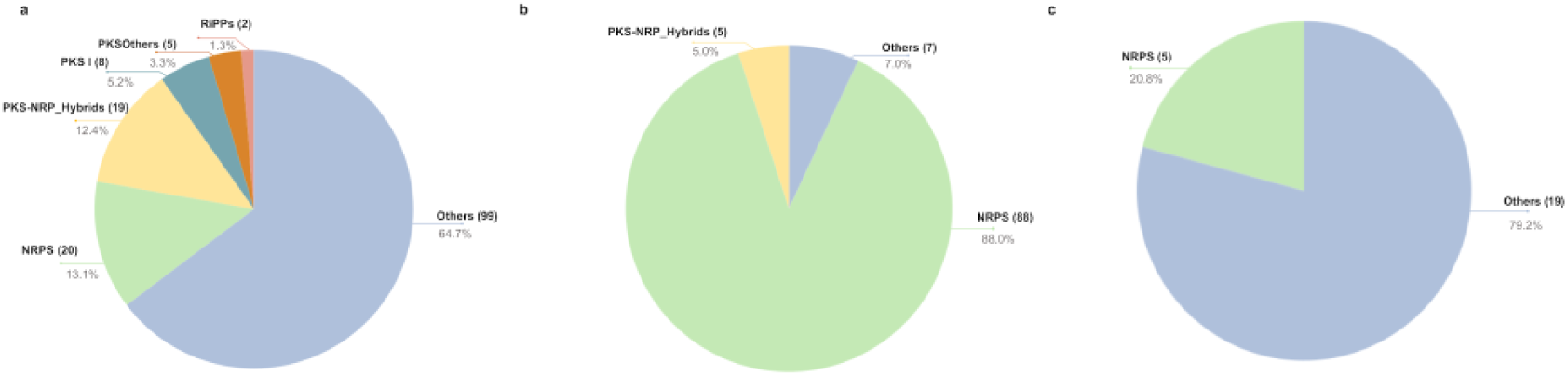
BGC per biosynthetic class distribution for potentially. a) benzoxazolinate, b) benzobactin, and c) ashimide-like BGCs highlights the diverse biosynthetic origins of clusters.

### Rapid and accurate screening of BGCs using a neural network

For screening rare clusters encoding potential benzoxazolinate-associated SMs, we employed a multilayer perceptron (MLP) neural network framework to develop a bioprospecting model that learns nucleotide features directly from BGC sequences. The model trained on 6-mer nucleotide frequencies and evaluated using an 80:20 holdout approach, achieved an average accuracy of 93.5%. Additional experiments using 10-fold cross-validation (CV), the model achieved 91.7% accuracy. Performance evaluation using class-wise metrics demonstrated reliable classification of both positive and negative classes. The classifier achieved F1-scores of 93% for the positive class and 94% for the negative class (Table S4).

Evaluation on an external validation dataset of 252 BGCs resulted in 97% accurate predictions. In addition, six experimentally characterized BGCs, which were not part of the training dataset, were classified correctly, supporting the utility of the framework on independent datasets. As a further proof-of-concept application, two metagenome-derived contigs were screened and predicted to contain benzoxazolinate-associated BGCs (Table S5).

The performance of the model can be attributed to the use of k-mer–based sequence features, which effectively capture local and compositional patterns in the sequences and do not require any gene annotation and predictions. The predictions of this model make it a suitable proof-of-concept prioritization tool for potential BGCs encoding benzoxazolinate-like SMs.

### Evolutionary history of hosts harboring benzoxazolinate-like BGCs

To assess the evolutionary relationship among genome mined bacterial hosts, gene trees for representative BGCs were constructed using protein sequences of XsbA, XsbC, and XvbB (Table S6). The representative phylogenetic trees of the XsbC and XsbA protein homologs derived from benzobactin and benzoxazolinate BGCs revealed genus-level clustering that highlights evolutionary conservation within the taxa (Fig. 5a-b, 6a-b). Similarly, the maximum likelihood tree based on the SHMT protein sequence is conserved within the taxa highlighted by genus-level clustering (Fig. S6). These results suggest that the core biosynthetic machinery is broadly conserved across phylogenetically related taxa.

**Figure 5:**
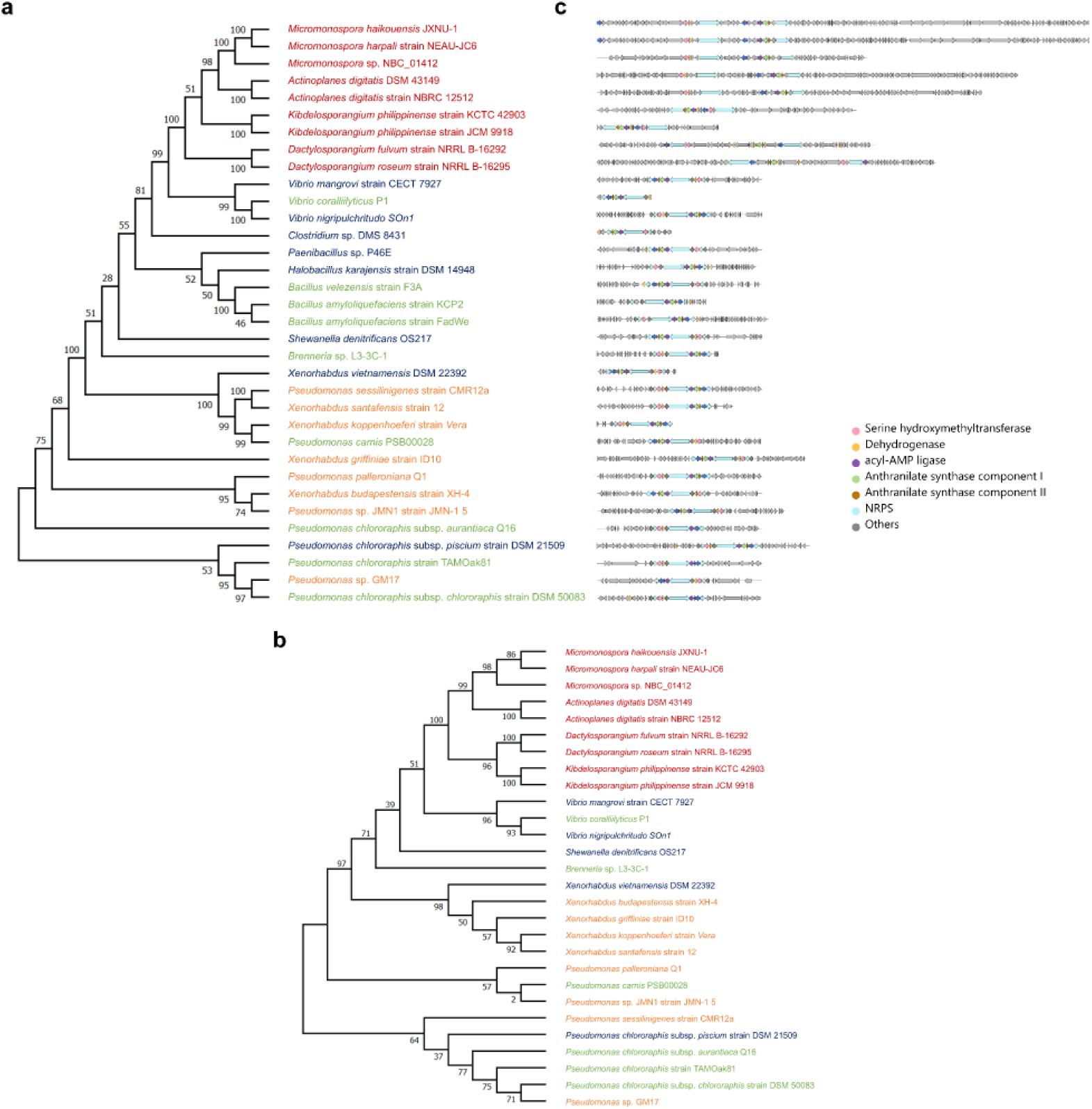
Phylogenetic profiling and gene architecture of benzobactin-like BGCs. The maximum-likelihood tree was constructed based on aligned a) acyl-AMP ligase b) dehydrogenase protein sequences derived from 34 representative BGCs associated with benzobactin-like compounds. Protein sequences were aligned using the ClustalW algorithm in MEGA11, and the phylogenetic tree with bootstrap support values shown at branch nodes. c) Corresponding BGC architectures (right) are shown as gene cluster maps, illustrating the conservation of core biosynthetic, regulatory, and transporter genes. The BGCs were annotated using antiSMASH and visualized by clinker. In panels a) and b) bacterial names in blue represent known species, dark red represent previously unreported genera, orange represent previously unreported species of known genera, and green represent previously unreported strains of known species.

**Figure 6:**
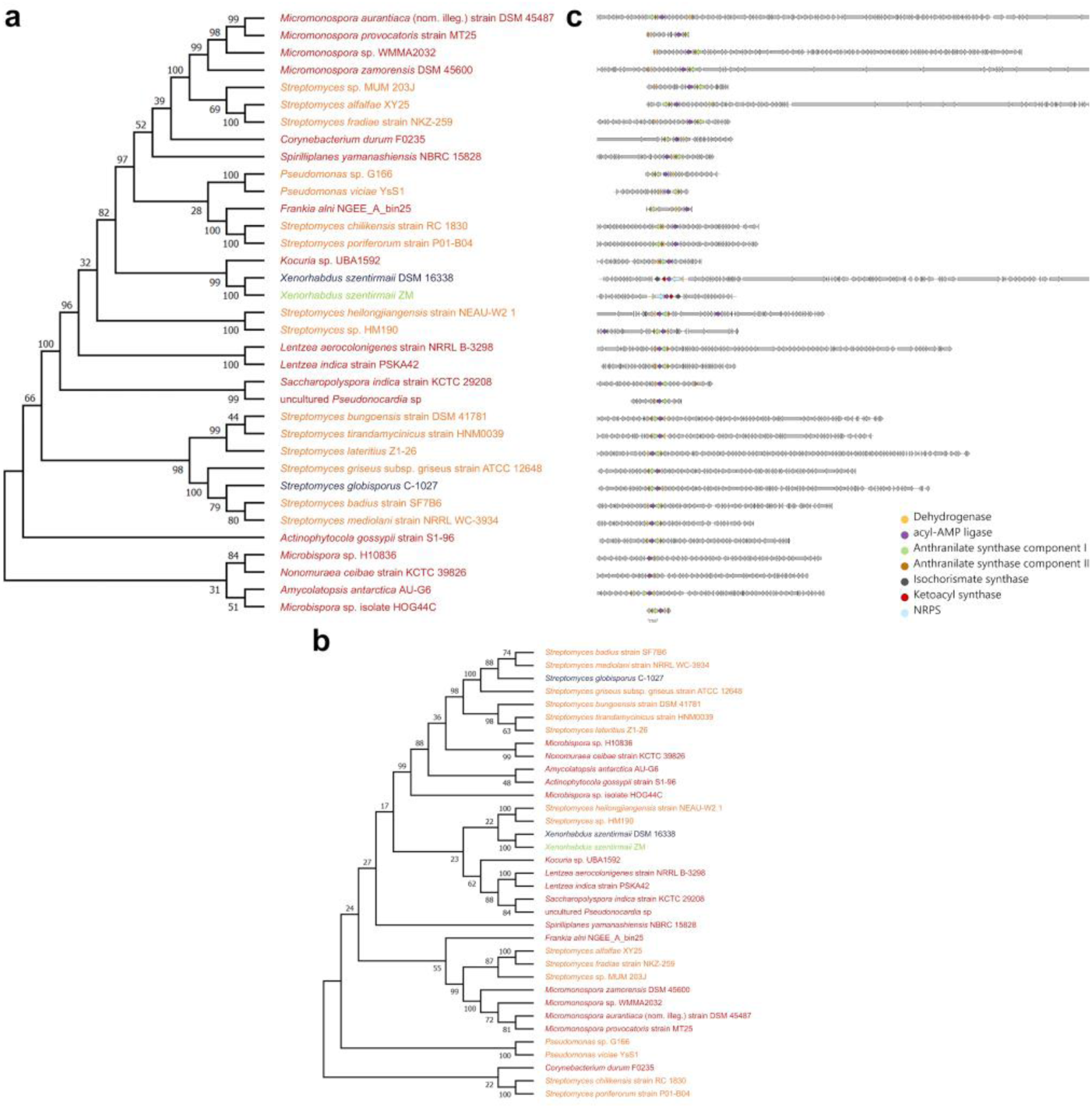
Phylogenetic profiling and gene architecture of benzoxazolinate-like BGCs. The maximum-likelihood phylogenetic tree was constructed based on aligned a) dehydrogenase and b) acyl-AMP ligase protein sequences derived from BGCs associated with benzoxazolinate-like compounds. c) Corresponding BGC architectures (right) are shown as gene cluster maps, illustrating the conservation of core biosynthetic, regulatory, and transporter genes. The BGCs were annotated using antiSMASH and visualized by clinker. In panels a) and b) bacterial names in blue represent known species, dark red represent previously unreported genera, orange represent previously unreported species of known genera, and green represent previously unreported strains of known species.

Further, the benzobactin representative clusters were also aligned and visualized using clinker (Gilchrist & Chooi, 2021), which highlights the conserved as well as rearranged architecture of core biosynthetic genes across diverse bacterial taxa, notably in previously unreported genera such as *Micromonospora*, *Actinoplanes*, *Kindelosporangium*, and *Dactylosporangium* (Fig. 5c). BGCs from these genera displayed a dispersed arrangement of core and accessory genes relative to other clusters, suggesting diversification in cluster architecture.

In the potential benzoxazolinate-producing bacterial hosts, the BGC architecture closely resembled the *sgc* cluster identified in *Streptomyces globisporus*. Each cluster contained only four core genes - dehydrogenase, acyl-AMP ligase, anthranilate synthase component I, and anthranilate synthase component II, as observed in *Pseudomonas*, *Micromonospora*, *Corynebacterium*, *Soirilliplanes*, *Frankia*, *Kocuria*, *Lentzea*, *Saccharopolyspora*, *Actinophytocola*, *Microbispora*, *Nonomuraea*, and *Amycolatopsis* genera (Fig. 6c). The maximum likelihood tree constructed from the representative BGCs encoding ashimides revealed conserved sequence homology within the bacterial genera, as well as conserved and rearranged BGC architecture across previously unreported genera (Fig. S7). These gene trees play an important role in identifying the events of gene expansion and subsequent diversification (Chevrette et al., 2021).

The phylogenetic analysis of major facilitator superfamily (MFS) transporters revealed clustering patterns among transporters of the same compound class, suggesting sequence conservation within these groups (Fig. S8). MFS transporters belong to multidrug (antibiotic) resistance (MDR) efflux pumps. These efflux pumps not only aid in conferring resistance to drugs and survival in hosts but also cellular detoxification of antimicrobial compounds (Piddock, 2006; Vela-Corcía et al., 2019). These observed clustering patterns may reflect functional specialization associated with the transport of structurally related metabolites, although this hypothesis requires further experimental validation.

### Predicted bioactive potential of genome mined BGCs

To predict the bioactivities of BGCs encoding potential benzoxazolinate-containing NPs, we used NPBdetect, a neural network-based model, to identify promising clusters with diverse bioactivities (Goyat et al., 2025). Among the 277 analyzed BGCs, 225 were predicted to be associated with antibacterial activity, followed by 84 with cytotoxic/antitumor activity, 48 with antifungal, and 11 with siderophore activities (Table S7). An UpSet plot (Conway et al., 2017) was generated using the prediction output to visualize the intersection of predicted bioactivities, revealing that of the 225 antibacterial predicted BGCs, 117 BGCs were predicted exclusively antibacterial, whereas 56 BGCs showed overlap of antibacterial and cytotoxic bioactivity predictions (Fig. S9).

To further strengthen confidence in predicted bioactivities, we used a machine learning model, the natural product function (NPF) tool, to predict antibacterial, antifungal, and antitumor/cytotoxic bioactivities (Walker & Clardy, 2021). NPF predictions revealed 252 BGCs predicted to be antibacterial and 84 BGCs predicted to be cytotoxic (Table S8). Overlap analysis of the predicted bioactive BGCs from both prediction tools resulted in the identification of 213 antibacterial and 41 cytotoxic BGCs (Fig. S10).

Out of the 529 genome mined BGCs, 32 were predicted to exhibit siderophore-like bioactivity, of which five encode benzoxazolinate-like and the remaining 27 correspond to benzobactin-like clusters (Table S9). The analysis of siderophore-associated domains within the predicted siderophore BGCs revealed the presence of a single TonB-dependent receptor across eight benzobactin BGCs, whereas four benzoxazolinate-like BGCs encoded both periplasmic binding proteins (PBP) and FecCD transporter domains and followed the antiSMASH nonribosomal independent (NI)-siderophore rule. Interestingly, four of the benzoxazolinate BGCs harboring both FecCD and PBP domains were clustered together, highlighting a conserved cluster and transporter organization (Table S9). The predicted siderophore and cytotoxic bioactivities highlight these BGCs as promising candidates for future experimental studies. Further functional characterization may confirm their dual bioactivities and provide insights into their biosynthetic potential.

### Openness metrics highlight gene composition diversity

To quantify compositional diversity within the cluster families, openness metric analysis was performed across the identified GCFs. According to established thresholds (*γ* < 0.3: closed; 0.3 − 0.6: intermediate; *γ* > 0.6: open) (Hyun et al., 2022; Paccagnella et al., 2025; Rajput et al., 2023), the majority of the GCFs lies in the closed to intermediate range, indicating a limited increase in gene composition diversity with the addition of related BGCs (Table S10). However, in FAM_02558 and FAM_02717, the pangenome metric values are 0.743 and 0.997, respectively, suggesting greater variability in gene composition among member clusters. The average unique BGC gamma value of 0.983 indicates modular gene reorganization. Overall, these findings indicate variable patterns of gene composition diversity across the identified GCFs.

### Identification of tailoring enzyme in genome mined BGCs

Across the 277 BGCs, tailoring enzyme annotations were obtained from antiSMASH v8, as this feature is not available in the earlier versions. Among these, 18 BGCs contained tailoring enzymes with corresponding annotations in the MITE database. The majority of these BGCs (n=14) harbored enzymes showing similarity to MITE0000068 (Oxidation), while two BGCs each associated with MITE0000077 (Methylation) and MITE0000104 (Hydroxylation) (Table S12). These sequence similarity based tailoring annotations suggest the potential for diverse tailoring reactions, which may contribute to structural and functional diversification of the encoded compounds.

### Distribution of candidate BGC hosts across rhizosphere samples

In the genome mined dataset, we identified two benzoxazolinate-like BGCs from metagenome assembled genomes (MAGs), including *Frankia alni* NGEE_A_bin25 and uncultured *Pseudonocardia sp.*, sourced from the SMC database. Similarity searches against ABC-HuMi and BGC Atlas detected homologs of core biosynthetic proteins, but no overlapping regions indicative of complete candidate BGCs were identified (Fig. S11-S12). As a pilot study, we analyzed the rhizosphere metagenomic samples (Zhou et al., 2025) to study the ecological distribution of genome mined bacterial hosts. Out of the 268 analyzed bacterial strains, 178 bacterial strains were identified to be abundant across the rhizosphere samples with relative abundance values ranging from 0 to 19.42. *Lentzea aerocolonigenes* strain NRRL B-3298 was identified as the most abundant bacterial strain in switchgrass samples, followed by *Pseudomonas sp.* MWU13-2100 and *Pseudomonas sp.* G2-4 (Fig. 7). These results provide an overview of the distribution of candidate BGC-associated hosts in rhizosphere microbiomes. Additionally, we were able to screen two benzoxazolinate-like clusters from two assembled metagenome contigs through our bioprospecting model (Table S5). Importantly, the recovery of benzoxazolinate-like clusters directly from metagenomic contigs highlights the utility of our screening framework and warrants identifying promising targets for downstream experimental characterization and natural product discovery.

**Figure 7:**
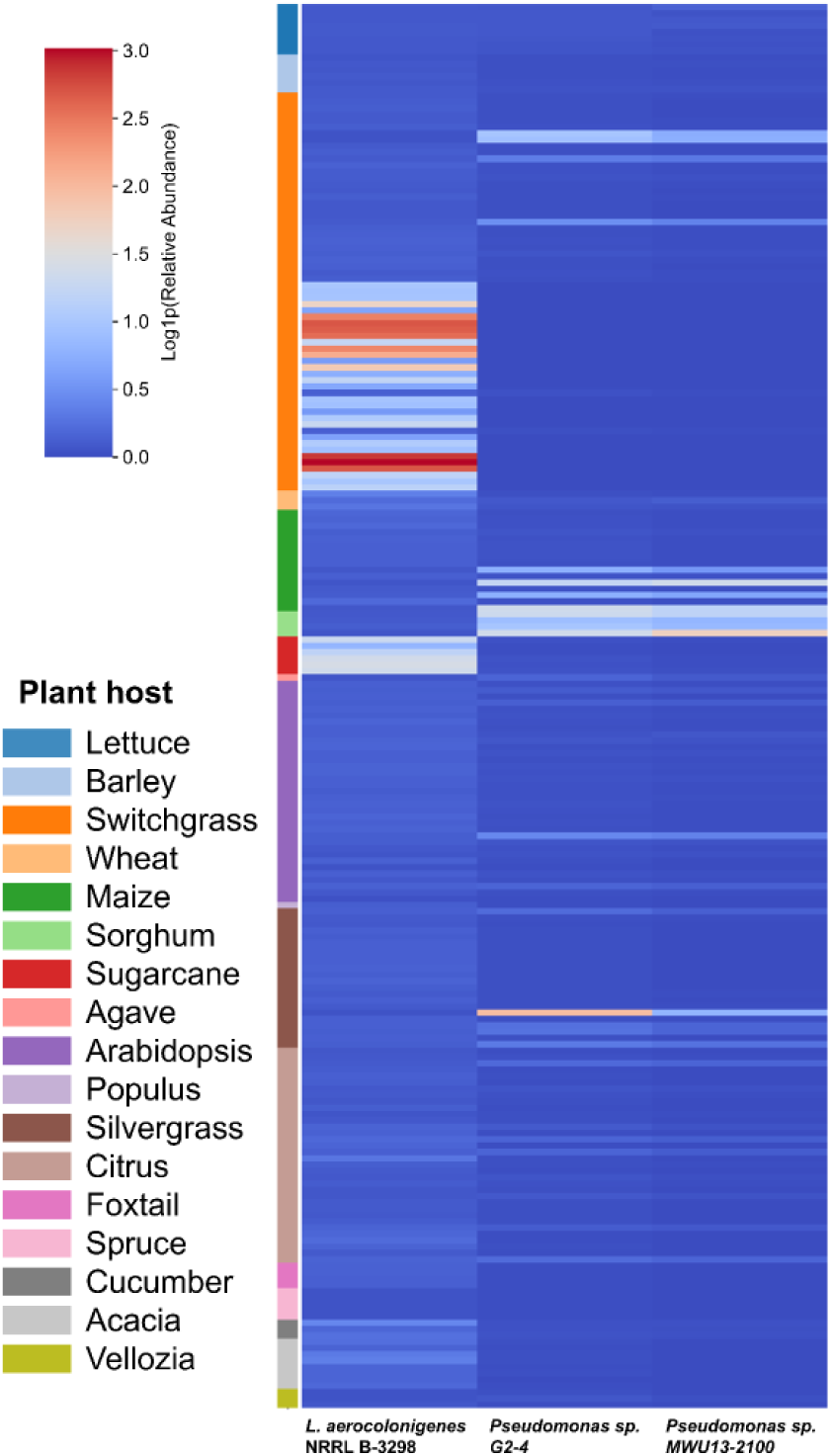
Relative abundance of genome mined benzoxazolinate-producing bacterial strains across rhizosphere samples. Relative abundance of top three most abundant bacterial strains across 222 analyzed rhizosphere samples highlights significant abundance of the *Lentzea aerocolonigenes* strain NRRL B-3298 across switchgrass, followed by *Pseudomonas sp.* MWU13-2100 across silver grass and *Pseudomonas sp*. G2-4 across sorghum.

### Future outlook, recommendations, and limitations

The bioactivity predictions and subsequent analysis of the transporter domain and antiSMASH siderophore rules highlight the siderophore potential associated with the identified clusters. Currently, there are no reports of the siderophore activity of benzoxazolinate-associated NPs. Experimental validation will therefore be essential to ascertain the role of these metabolites in iron acquisition and to elucidate their ecological and physiological roles. Future investigations should also experimentally validate the substrate specificity and transport kinetics of benzoxazolinate-associated MFS transporters to test the hypothesis that their evolutionary clustering mirrors chemical class–dependent detoxification and self-resistance mechanisms in the producing host.

The distribution of benzoxazolinate-associated BGCs across biosynthetic classes, including NRPS, PKS, other, and PKS-NRP_hybrids, highlights the limitations of the current biosynthetic classification framework. Given the presence of a conserved gene cassette involved in benzoxazolinate core assembly (including acyl-AMP ligase, dehydrogenase, ADIC synthase) and its derivatives, we propose a preliminary antiSMASH detection rule, based on profile HMMs (pHMMs) of the core genes for improved identification of benzoxazolinate-associated BGCs (Fig. S13) (Reitz et al., 2026). These refinements would help distinguish these rare bis-heterocyclic scaffolds from canonical NRPS- or PKS-derived clusters and prevent their misclassifications into broad, functionally diverse categories.

To date, the MIBiG repository contains only two experimentally validated BGCs associated with benzoxazolinate-derived metabolites: C-1027 and ashimide. The *xsb* BGC from *X. szentirmaii*, alongside the *pbz* cluster from *P. chlororaphis* (Y. Shi et al., 2022) and *xvb* cluster from *X. vietnamensis* (Y.-M. Shi et al., 2022) encoding benzoxazolinate and benzobactin, represent additional validated candidates that warrant inclusion in the MIBiG database next release.

A key limitation of this study is that the WGS dataset was searched in a taxon-specific manner, focusing only on selected genera and species, which may have led to the omission of additional benzoxazolinate-like BGCs present in other microbial taxa. Additionally, the developed bioprospecting MLP model is primarily designed as a proof-of-concept prioritization tool and does not provide functional annotation of the screened cluster. Furthermore, as the model is trained on in silico-predicted data, it needs to be benchmarked against experimentally validated BGCs. However, at present there are only six experimentally validated BGCs encoding benzoxazolinate containing NPs. Finally, the functional characterization and heterologous expression of the identified BGCs will be essential to explore their full biosynthetic potential.

### Conclusion

Through targeted BGC based genome mining, we identified 277 candidate BGCs that potentially encode NPs containing benzoxazolinate moiety from diverse bacterial hosts, including benzoxazolinate, benzobactin, and ashimide-like SMs. The candidate BGCs were distributed across diverse bacterial taxa, with maximum abundance observed in members of *Streptomycetaceae* and *Pseudomonadaceae*. These findings substantially expand the known biosynthetic repertoire beyond previously identified bacterial taxa. The clustering of identified BGCs with known MIBiG BGCs highlights conserved cluster architecture, whereas distinct GCFs and singletons suggest unexplored diversity and rearranged BGC architecture. Openness metric analysis across the identified GCFs revealed variable patterns of compositional diversity. The bioactivity predictions of the identified BGCs highlight the diverse bioactive potential associated with candidate BGCs, with predominance of antibacterial and cytotoxic BGCs, thereby warranting future experimental characterization. The rhizosphere microbiome survey revealed a notable abundance of genome mined bacterial strains producing benzoxazolinate-like NPs, suggesting the presence of potential clusters within the diverse microbiome. Furthermore, we developed a bioprospecting model based on MLP to rapidly screen BGCs for the potential of benzoxazolinate-containing NPs with high accuracy. This study also highlights the identification of benzoxazolinate-like producing microbes with dispersed BGCs and the need to devise new algorithms to annotate such rare dispersed BGCs as a single unit. The findings not only expand the catalog of bioactive SMs but also provide promising leads for future drug discovery, particularly in the context of antimicrobial resistance and cancer therapeutics.

## Materials and Methods

### Genomic data collection

To identify the potential bacterial hosts that produce benzoxazolinate-containing SMs, the protein sequences of *xsbA, xsbC, xpzC, and xvbB*, which are signature genes involved in benzoxazolinate and benzobactin biosynthetic pathways, were used as queries in tblastn searches. Searches were carried out against the complete prokaryotic genome database (comprising 144,111 sequences) (*National Center for Biotechnology Information*, accessed in March 2025), the WGS dataset of selected bacterial taxa (Table S12), and the BGCs available in the SMC database (containing 10,971,814 BGCs) (Udwary et al., 2025). Additionally, comprehensive genome mining was conducted by analyzing the NCBI nr-protein database and screening for protein homologs using blastp. Further BiG-FAM database (Kautsar et al., 2021) was queried using the antiSMASH job ID for the benzobactin reference BGC (*X. vietnamensis* DSM 22392). The resulting GCF was analysed to identify BGCs associated with the benzobactin biosynthesis pathway. Venny v2.1.0 (Oliveros, 2007) was used to analyze the common significant hits obtained from each query protein. Bacterial genomes containing combinations of all four proteins of interest or of three proteins of interest were selected. The selected bacterial genomes were downloaded from the NCBI nucleotide database using the Batch Entrez utility (https://www.ncbi.nlm.nih.gov/sites/batchentrez).

### Genome mining for identification of target BGCs

The nucleotide sequences for the selected bacterial genomes and gene clusters shortlisted from the SMC database were analyzed for BGCs encoding benzoxazolinate-containing SMs using the command-line version of antiSMASH v7.1 (Blin et al., 2023). Parameters were as follows: – minlength 5000, –cb-knownclusters, –cb-general, –cb-subclusters, –asf, –pfam2go, –smcog-trees, –genefinding-tool prodigal, –fullhmmer.

The GenBank (gbk) files of the antiSMASH-annotated BGCs were obtained and used to create a nucleotide BLAST database using a local installation of the NCBI BLAST package (version 2.16.0+). The resulting database was then queried against the amino acid sequence of XsbA, XsbC, XpzC, and XvbB using the tBLASTn algorithm. Venny v2.1.0 was used to analyze the common significant hits obtained from each query protein. This targeted search strategy allowed the investigation of candidate benzoxazolinate-associated BGCs, thereby reducing background noise. The BGCs containing combinations of all four proteins of interest or a combination of three proteins of interest were subjected to deduplication using Dedupe v39.33 of the BBTools package (https://jgi.doe.gov/data-and-tools/bbtools/). The resultant non-redundant BGCs were selected for further analysis. Isolation source information for the BGCs was obtained from NCBI, and the global distribution map was constructed using geographic coordinates curated in a Google Spreadsheet.

### Sequence similarity network construction

The filtered BGCs were obtained and processed locally for clustering analysis using Biosynthetic Genes Similarity Clustering and Prospecting Engine (BiG-SCAPE) (Navarro-Muñoz et al., 2020) with the Minimum Information about a Biosynthetic Gene cluster (MIBiG) database v3.1 (Terlouw et al., 2023) as reference (using ‘–mibig’ parameter). The BGCs were grouped into GCFs, and a sequence similarity network of BGCs was constructed using a score cutoff (c=0.3). The GCFs clustering the antiSMASH-processed BGCs and MIBiG representative BGCs were considered as known GCFs. Based on their sequence similarity networks, the BGCs were classified into three different potential compound classes: benzoxazolinate, benzobactin, and ashimides.

### Bioprospecting using neural network

In addition to the traditional bioprospecting method, including sequence-based database searches, we developed a MLP neural network to screen BGCs encoding benzoxazolinate-containing SMs. This model uses the nucleotide sequences to screen BGCs for their biosynthetic potential. The MLP neural network was implemented using PyTorch and consisted of three fully connected layers with 128 and 64 hidden units. Rectified linear unit (ReLU) activation functions were applied after each hidden layer. In the literature, only six BGCs have been experimentally characterized for their biosynthetic potential to encode benzoxazolinate-containing SMs, limiting their use in training the model. As a result, we have used an in silico predicted genome mined BGCs dataset as a silver-standard set to train the model (Wagholikar et al., 2020). 271 BGCs from the genome mined dataset and 269 BGCs from the MIBiG were taken as positive and negative datasets to train the model. From the nucleotide sequences, 4096 6-mer features were computed using a custom Python script, and the resulting values were organized into a k-mer feature matrix. The dataset was split into training and testing sets using an 80:20 ratio, and 10-fold CV was performed. The 252 (apart from 277 BGCs) from the 529 genome mined dataset (Table S13) and the six experimentally validated BGCs, which were not part of the training and test set, were used as an external validation dataset.

### Phylogenetic analysis

For the three compound classes, the phylogenetic trees were constructed using protein sequences from representative BGCs (Table S6) that produced significant alignments with XsbA, XsbC, and XvbB. The sequences were aligned using the ClustalW algorithm (default parameters), and maximum likelihood trees were constructed with Molecular Evolutionary Genetics Analysis (MEGA11), employing 1,000 bootstrap replicates to assess the statistical support for each branch (Tamura et al., 2021).

A comprehensive search for MFS transporters was conducted across 277 identified BGCs. A Pfam-derived hidden Markov model (HMM) was used to query protein sequences using hmmsearch. This analysis led to the identification of 574 MFS proteins distributed across 259 BGCs. The resulting phylogenetic tree was exported in Newick format and visualized in iTOL (Letunic & Bork, 2024) to examine clustering patterns among MFS proteins originating from distinct BGCs.

### Bioactivity predictions

To predict the biological activities of the genome mined BGCs, NPBdetect, a neural network model, was utilized (Goyat et al., 2025). NPBdetect can predict multiple bioactivities of SMs based on their BGC sequences, including antibacterial, antifungal, cytotoxic/antitumor, siderophore, antiviral, antiprotozoal, inhibitor, and surfactant activities. The gbk files generated from antiSMASH were used as input to predict these bioactivities. To increase the reliance on predicted bioactivities, we additionally utilized the NPF tool (Walker & Clardy, 2021) to predict antibacterial, cytotoxic/antitumor, and antifungal bioactivities, and compare the predicted bioactivities with those obtained from NPBdetect.

Additionally, the remaining 252 clusters from the complete 529 BGCs dataset (Table S13) were screened for bioactivity predictions, with a primary focus on siderophore bioactivity. The predicted siderophore-positive BGCs and associated bacterial genomes were further analyzed for the presence of siderophore-associated transporter domains, namely FecCD, PBP, and TonB (Crits-Christoph et al., 2021; Reitz & Medema, 2022) using HMMER v3.3.2 (Potter et al., 2018). The predicted BGCs were examined against antiSMASH siderophore rules to identify biosynthetic signatures corresponding to recognized siderophore structural classes, followed by the BiG-SCAPE clustering.

### Openness metrics calculation

To assess the degree of compositional variability within the GCFs, we employed PanBGC to quantify openness metrics adapted specifically for BGCs (Paccagnella et al., 2025). For this analysis, we selected GCFs from BiG-SCAPE that contained MIBiG references for C-1027 and ashimides, along with GCFs with experimentally validated BGCs encoding benzoxazolinate and benzobactin. We also included the top GCFs for each biosynthetic class in the analysis. These GCFs were processed using the command line, and their data were visualized by uploading the visualization.json files to the PanBGC web server.

### Tailoring enzyme profiles

To identify the tailoring enzymes associated with the genome mined BGCs, we parsed the antiSMASH 8 (Blin et al., 2025) generated “index.html” files. Information on tailoring enzymes annotations with >60% similarity with the MITE database was extracted. Information on their corresponding locus, enzyme class, sub-functions, descriptions, and associated MITE IDs were also retrieved. These annotations were used to analyze the distribution of tailoring enzymes among the identified BGCs.

### Microbiome abundance profiling of genome mined bacterial strains

To identify the candidate benzoxazolinate-like BGCs in MAGs, we screened publicly available datasets including BGC Atlas (Bağcı et al., 2025) and ABC-HuMi (Hirsch et al., 2024). Core biosynthetic proteins (XsbA, XsbC, XpzC, and XvbB) were used as probes to search putative BGCs from curated databases, followed by overlap analysis to identify common regions with candidate clusters.

Further to assess the ecological distribution of genome mined bacterial strains across plant-associated microbiomes, we analyzed a total of 222 publicly available rhizosphere metagenome datasets (Zhou et al., 2025). Taxonomic classification was performed with Kraken2 (Lu et al., 2022) using a custom-built database of genome mined bacterial strains and a kraken standard database. Kraken2 report outputs were parsed to extract the relative abundance of target species across all samples, and the results were compiled into a species-by-sample abundance matrix.

## Data and code availability

The data presented in the paper and the codes of this study are available for download via the Google Drive link: https://drive.google.com/drive/folders/1cowVh4GwY408-TMZrmiYPkE1S6cPmShr?usp=sharing

All phylogenetic trees are interactive and can be accessed online on iTOL: (https://itol.embl.de/shared/1xPwOAB1weRj5).

## Acknowledgments

We thank the BRIC-National Agri-Food and Biomanufacturing Institute (NABI) for research facilities and supporting high-performance computing. We acknowledge the National Supercomputing Mission (NSM) for providing computing resources of PARAM Smriti at BRIC-NABI, Mohali, which is implemented by C-DAC and supported by the Ministry of Electronics and Information Technology (MeitY), Department of Biotechnology (DBT), and Department of Science and Technology (DST), Government of India (GOI). We acknowledge the assistance from the NCBI support desk for data retrieval. SM and SP would like to acknowledge Prof. Nadine Ziemert for her valuable comments and suggestions to improve the manuscript. We sincerely thank Martina Adamek for her valuable insights and thoughtful suggestions regarding the interpretation of results. SP is thankful to the Regional Center of Biotechnology (RCB), DBT, GOI, for undertaking PhD registration. SM would like to acknowledge NABI Core funding for the execution of this project. SM, AC, and LS acknowledge the financial support provided by the Department of Biotechnology (DBT), Ministry of Science & Technology, Government of India, under the project grant number [BT/PR54729/BSA/33/250/2024] and project BRIC-NABI Biomanufacturing BioFoundry “Agri-Food Bio-Mug” [BT/TEMP/24019/BMH-01/24].

## Additional files

Supplementary File Supplementary Tables

## Additional information

### Author’s contribution

SM and SP conceptualized and designed the study. SP collected the data and did the formal bioinformatic analysis. BK, LR, AC, IS, and MS assisted in data analysis. LS and SP developed the bioprospecting MLP model. SP and SM analyzed and finalized the results. SP, BK, LR, AC, IS, and MS prepared the manuscript. DS and VC critically reviewed the manuscript and provided valuable comments and suggestions for improving the draft. SM supervised and reviewed the manuscript.

