## Supplementary File for "Microbial bioprospecting for benzoxazolinate-like molecules: unleashing the potential of genome mining"

**List of supplementary figures**

Figure S1: Venn diagram displaying an overlap of 629 WGS sequences. This led to the identification of 37 WGS sequences containing combinations of the XsbA, XsbC, XpzC, and XvbB proteins.

Figure S2: Venn diagram displaying an overlap of SMC database-mined BGCs. This led to the identification of 164 BGCs containing a combination of the proteins XsbA, XsbC, and XvbB, and 241 BGCs containing a combination of XsbA and XsbC.

Figure S3: Venn diagram showing an overlap of 659 significant BGCs leading to the identification of 83 BGCs containing a combination of all four or three proteins of interest.

Figure S4: Workflow illustrating the collection, genome mining, and deduplication of genomic datasets used to identify 277 non-redundant BGCs with the potential to encode benzoxazolinate-containing SMs.

Figure S5: Global geographic distribution of 214 genome mined bacterial hosts encoding benzoxazolinate-associated clusters. Country metadata were unavailable for the remaining 63 identified hosts.

Figure S6: Phylogenetic tree of SHMT proteins identified in representative genome mined bacterial hosts with potential benzobactin-like BGCs.

Figure S7: Phylogenetic tree of a) dehydrogenase, c) acyl-AMP ligase, and d) SHMT proteins identified in representative genome mined bacterial hosts with potential ashimides-like BGCs. b) Corresponding BGC architectures (right) are shown as gene cluster maps, illustrating the conservation of core biosynthetic, regulatory, and transporter genes. The BGCs were annotated using antiSMASH and visualised by Clinker. Microbes in blue represent known species, Dark red represent novel genus, Orange represent novel species of known genus, and Green represent novel strain of known species.

Figure S8: The phylogenetic tree of 574 protein sequences of MFS transporters of 259 BGCs, highlighting clustering of proteins belonging to the same compound class. The green color clade represents benzobactin, red represents benzoxazolinate, purple represents C-1027, and blue represents ashimides-like BGCs.

Figure S9: Upset plot showing intersections of predicted bioactivities among BGCs. Vertical bars represent shared bioactivity combinations, while horizontal bars indicate the total number of BGCs per bioactivity class.

Figure S10: Venn diagrams showing overlap of BGCs predicted to be a) antibacterial and b) cytotoxic/antitumor by two bioactivity prediction tools, NPF and NPBdetect, strengthening the confidence in predicted bioactivities.

Figure S11: Screening ABC-HuMi database for putative benzoxazolinate-like BGCs: overlap analysis of core biosynthetic protein homologs of benzoxazolinate-like BGCs identified in MAGs sourced from the ABC-HuMi database revealed no overlapping metagenomic regions containing the required combination of protein homologs, indicating the absence of clusters in the analysed MAGs.

Figure S12: Screening BGC atlas database for putative benzoxazolinate-like BGCs: overlap analysis of core biosynthetic protein homologs of benzoxazolinate-like BGCs identified in MAGs sourced from the BGC Atlas database revealed no overlapping metagenomic regions containing the required combination of protein homologs, indicating the absence of clusters in the analysed MAGs.

Figure S13: Profile HMMs for proposed antiSMASH detection rules for benzoxazolinate-associated clusters.

**List of Supplementary Tables**

Table S1: Potential benzobactin BGCs belonging to GCF_13705 mined from the BiG-FAM database

Table S2: comprehensive list of 529 genome mined putative BGCs encoding benzoxazolinate-containing SMs

Table S3: List of putative 277 BGCs with microbial host information, potential compound encoded, novelty, genomic information, and isolation source information

Table S4: Performance of MLP NN model on positive and negative training sets

Table S5: Performance of MLP NN model on independent test set

Table S6: List of representative BGCs belonging to different compound classes for the phylogenetic analysis

Table S7: NPBdetect bioactivity predictions of 277 putative genome mined BGCs

Table S8: NPF bioactivity predictions of 277 putative genome mined BGCs

Table S9: List of 32 BGCs predicted to be siderophore positive mined from 529 BGCs and their siderophore-associated domains

Table S10: Openness metrics calculated for the GCFs obtained from the BiG-SCAPE clustering of genome mined BGCs

Table S11: antiSMASH annotated tailoring enzyme profiles of predicted genes of genome mined BGCs

Table S12: List of bacterial taxa selected for tblastn_vdb analysis

Table S13: 252 redundant BGCs from the 529 genome mined BGCs dataset

******

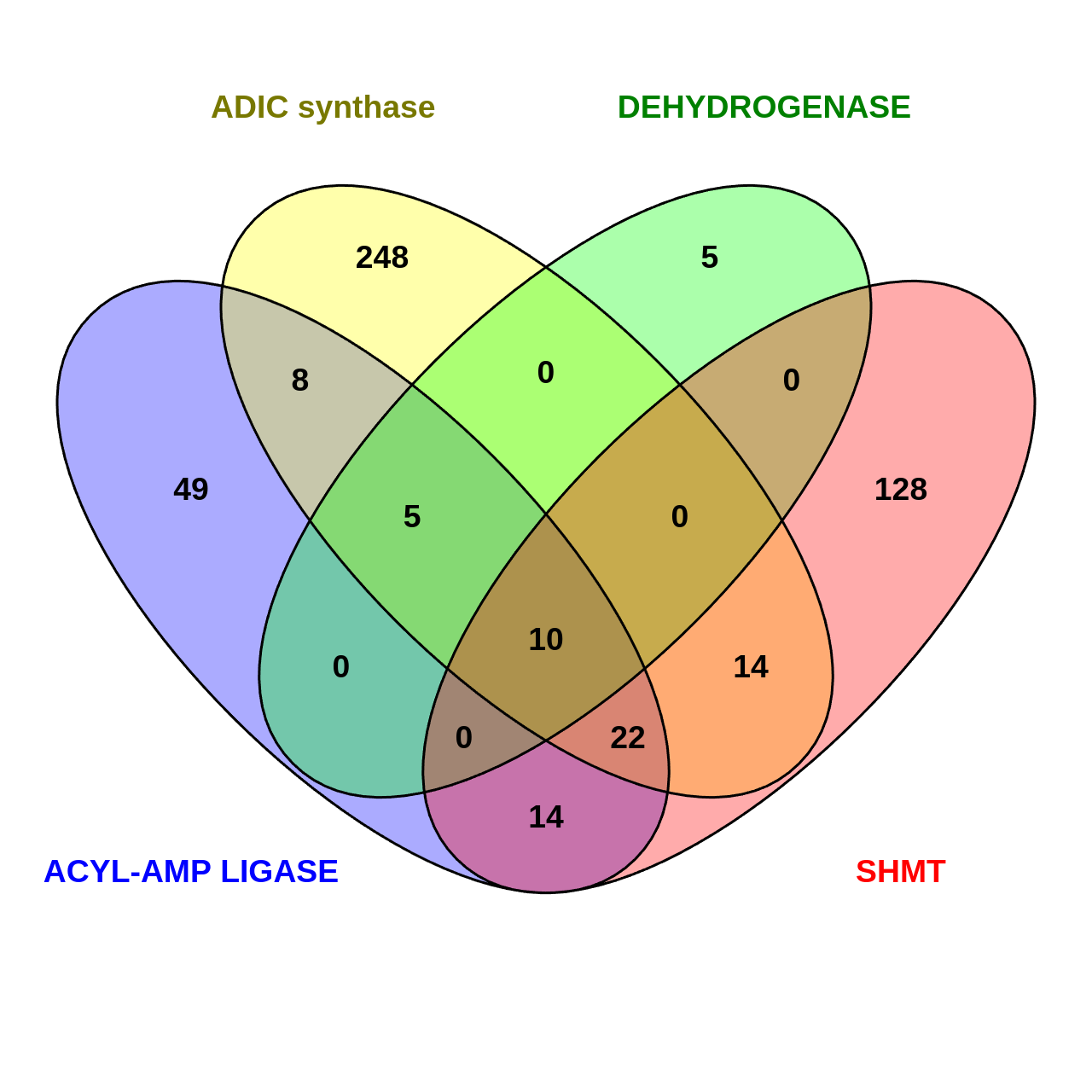

Figure S1: Venn diagram displaying an overlap of 629 WGS sequences. This led to the identification of 37 WGS sequences containing combinations of the XsbA, XsbC, XpzC, and XvbB proteins.

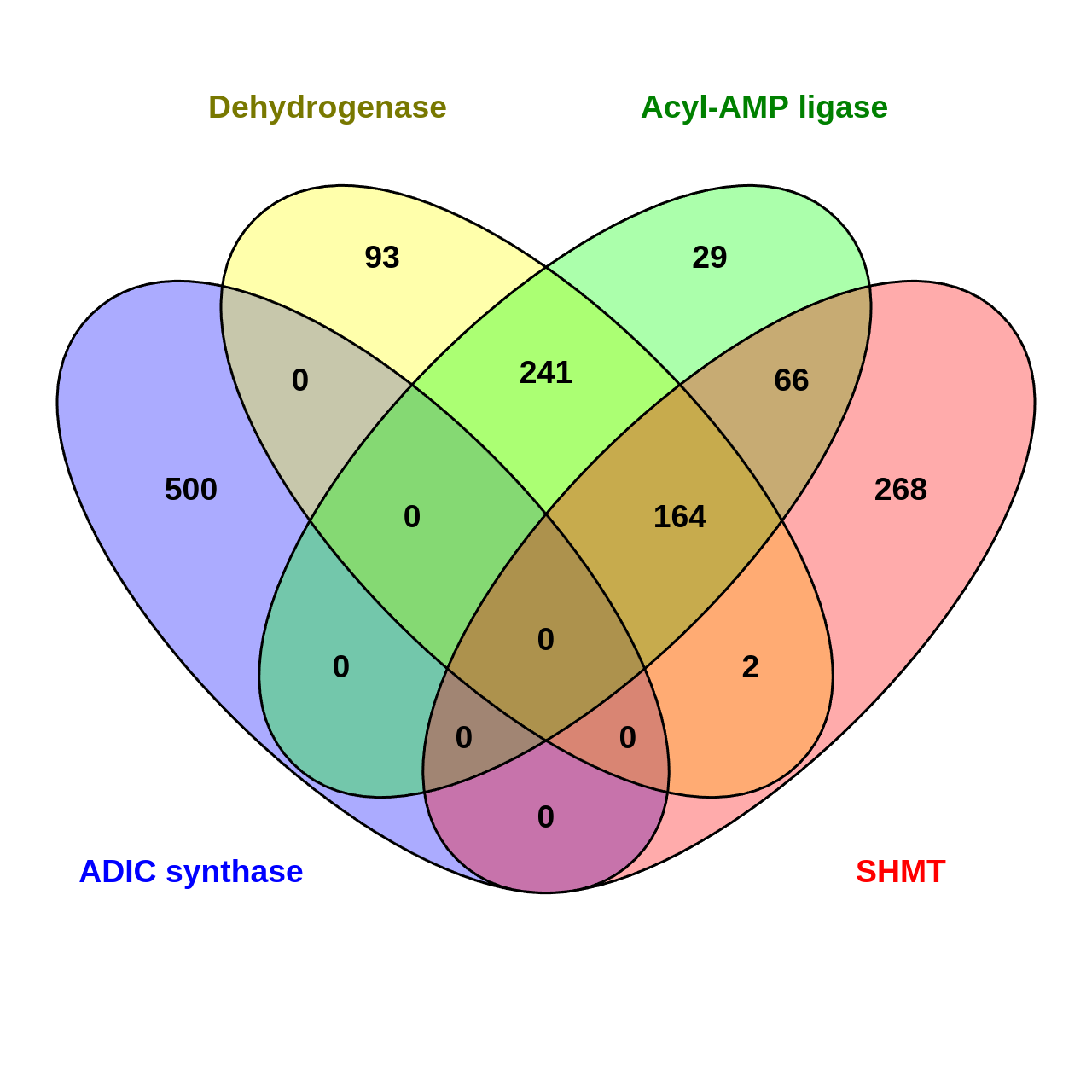

Figure S2: Venn diagram displaying an overlap of SMC database-mined BGCs. This led to the identification of 164 BGCs containing a combination of the proteins XsbA, XsbC, and XvbB, and 241 BGCs containing a combination of XsbA and XsbC.

.

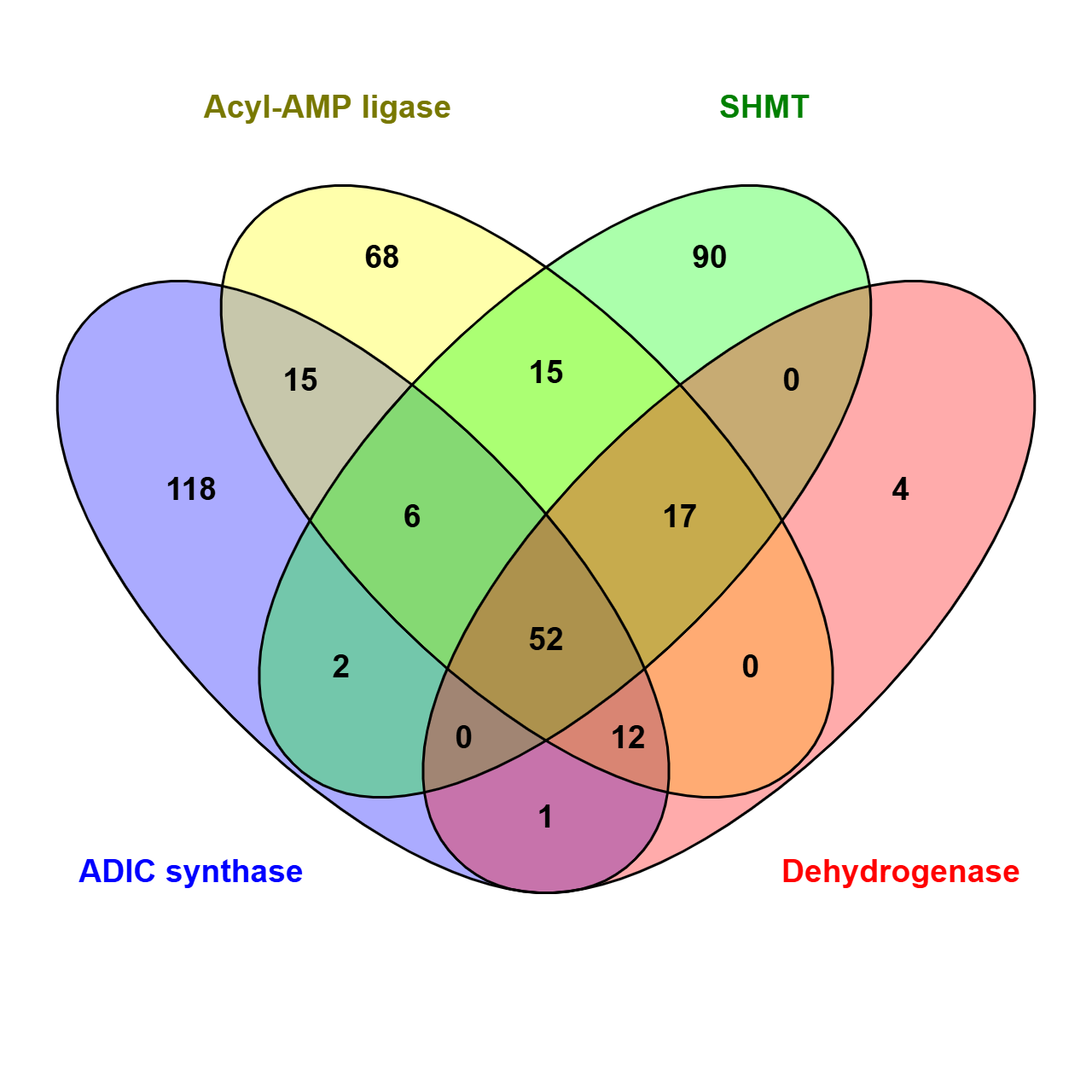

Figure S3: Venn diagram showing an overlap of 659 significant BGCs leading to the identification of 83 BGCs containing a combination of all four or three proteins of interest

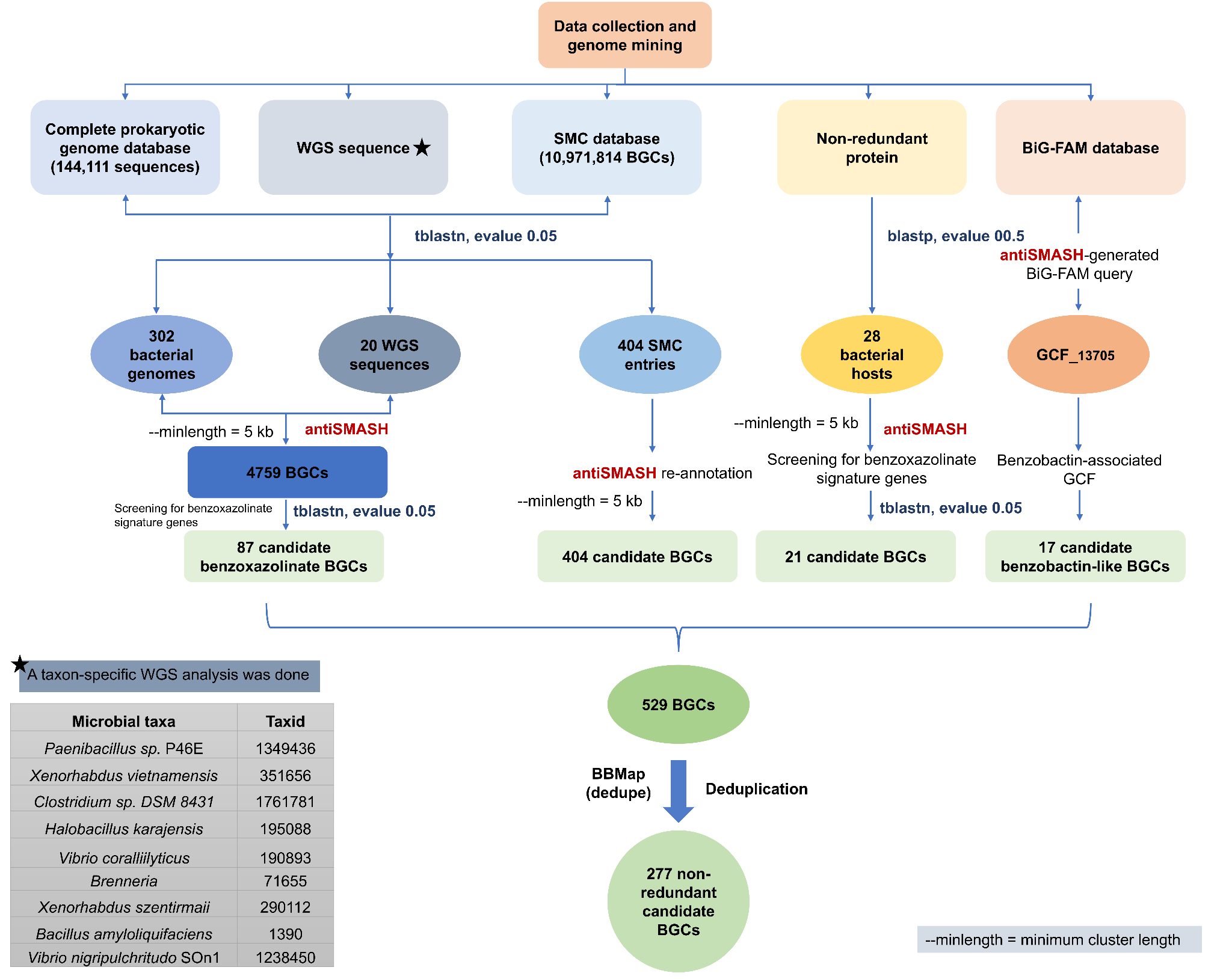

Figure S4: Workflow illustrating the collection, genome mining, and deduplication of genomic datasets used to identify 277 non-redundant BGCs with the potential to encode benzoxazolinate-containing SMs.

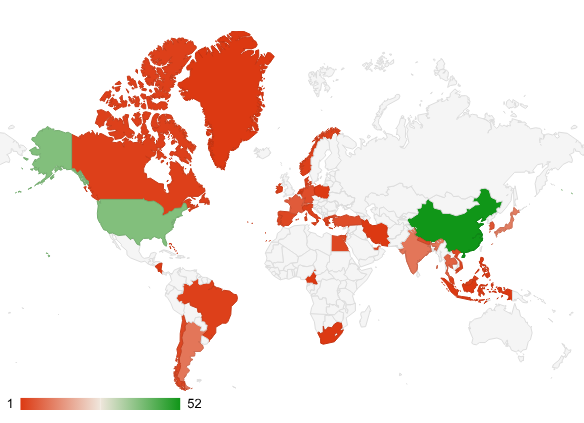

Figure S5: Global geographic distribution of 214 genome mined bacterial hosts encoding benzoxazolinate-associated clusters. Country metadata were unavailable for the remaining 63 identified hosts.

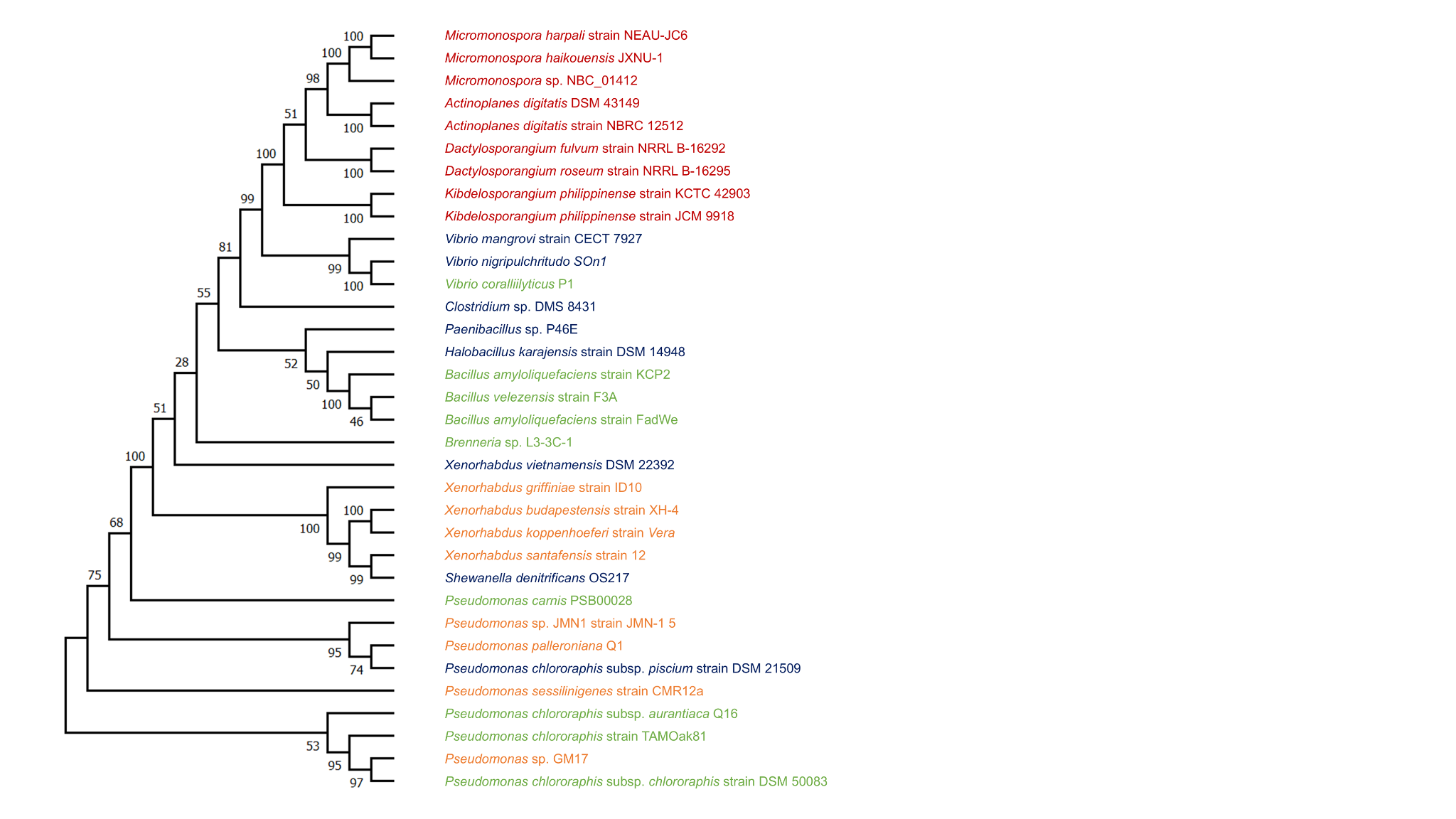

Figure S6: Phylogenetic tree of SHMT proteins identified in representative genome mined bacterial hosts with potential benzobactin-like BGCs

**iTOL**: <https://itol.embl.de/shared/1xPwOAB1weRj5>

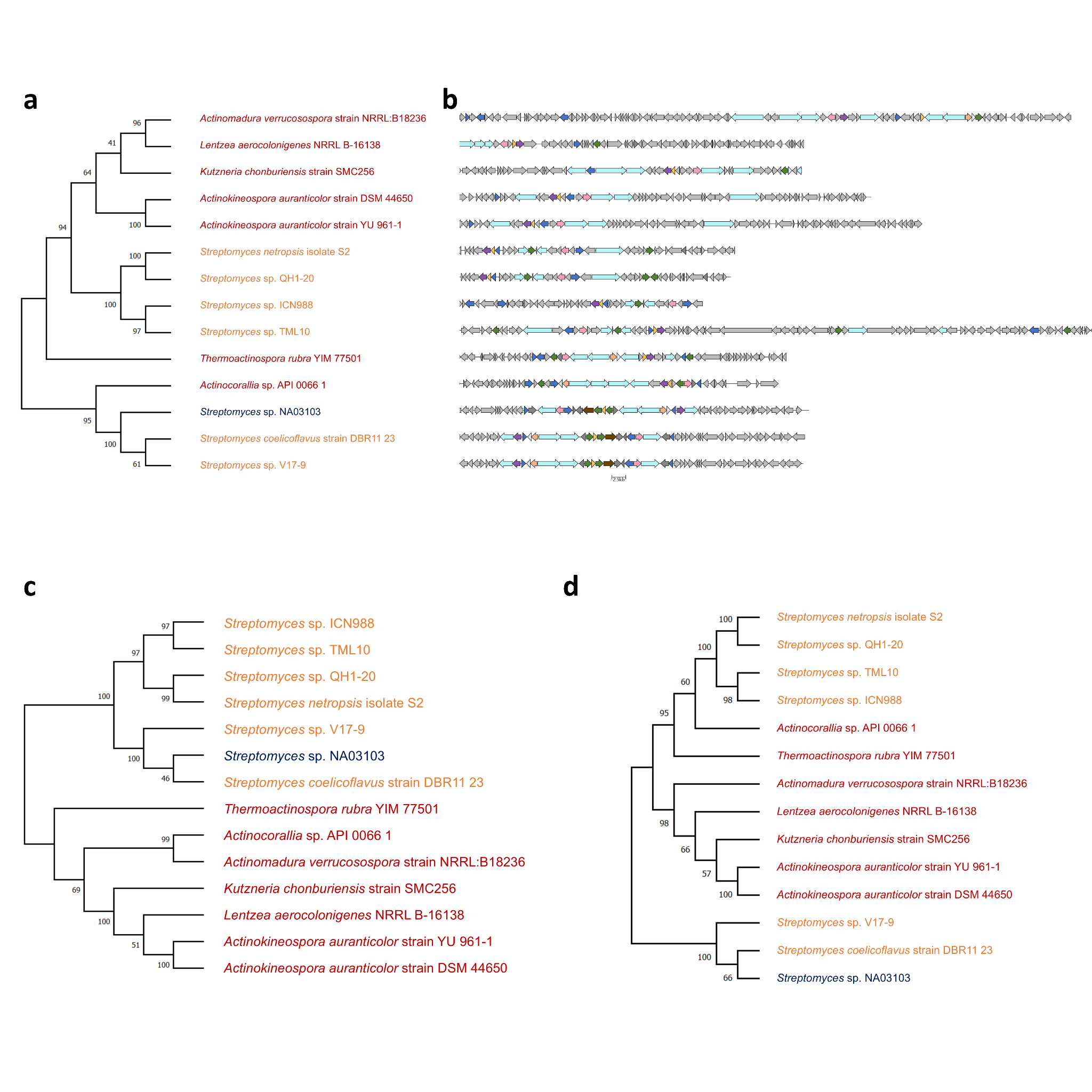

Figure S7: Phylogenetic tree of a) dehydrogenase, c) acyl-AMP ligase, and d) SHMT proteins identified in representative genome mined bacterial hosts with potential ashimides-like BGCs. b) Corresponding BGC architectures (right) are shown as gene cluster maps, illustrating the conservation of core biosynthetic, regulatory, and transporter genes. The BGCs were annotated using antiSMASH and visualised by Clinker. Microbes in blue represent known species, Dark red represent novel genus, Orange represent novel species of known genus, and Green represent novel strain of known species.

iToL link: <https://itol.embl.de/shared/1xPwOAB1weRj5>

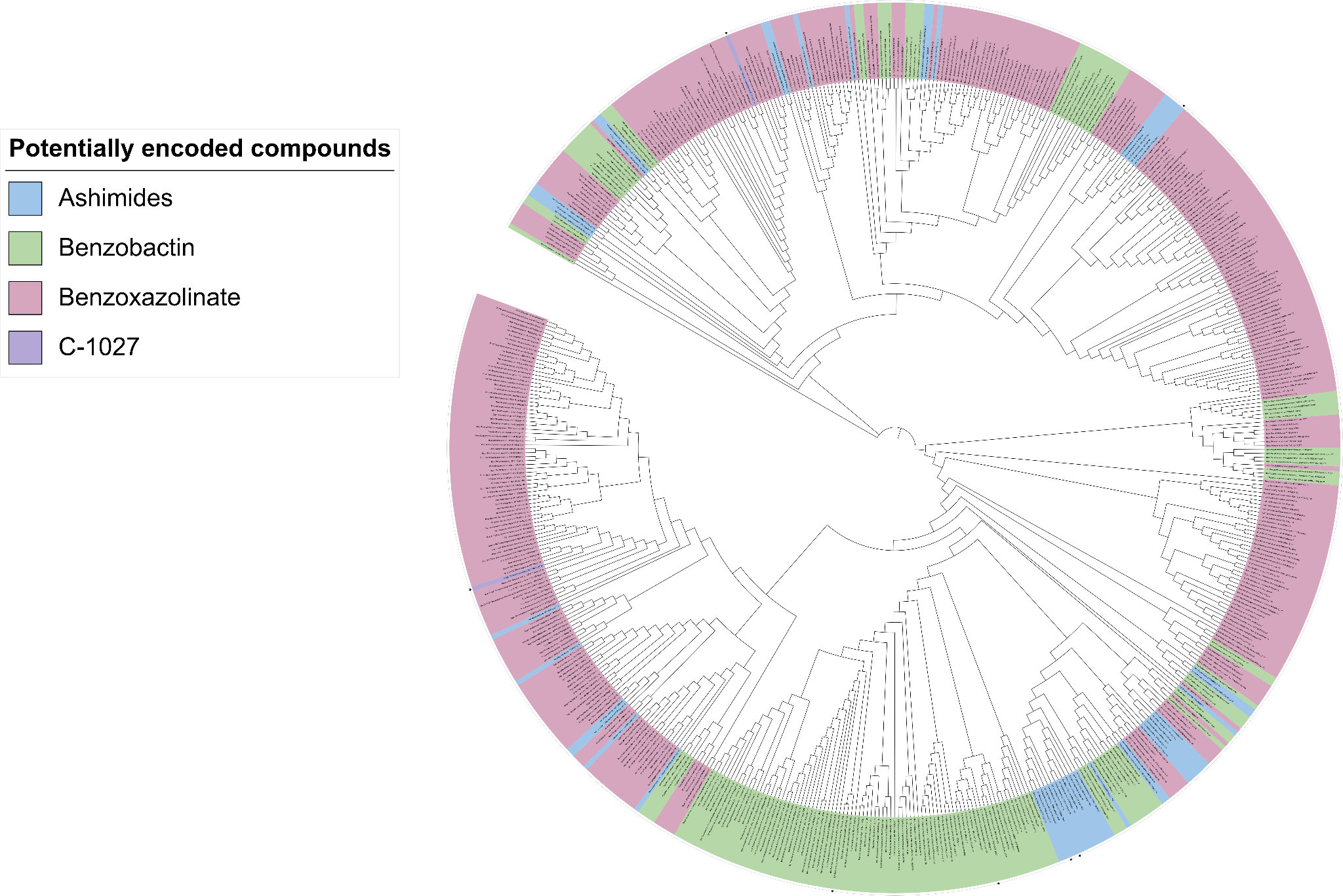

Figure S8: The phylogenetic tree of 574 protein sequences of MFS transporters of 259 BGCs, highlighting clustering of proteins belonging to the same compound class. The green colour clade represents benzobactin, red represents benzoxazolinate, purple represents C-1027, and blue represents ashimides-like BGCs.

iToL link: <https://itol.embl.de/shared/1xPwOAB1weRj5>

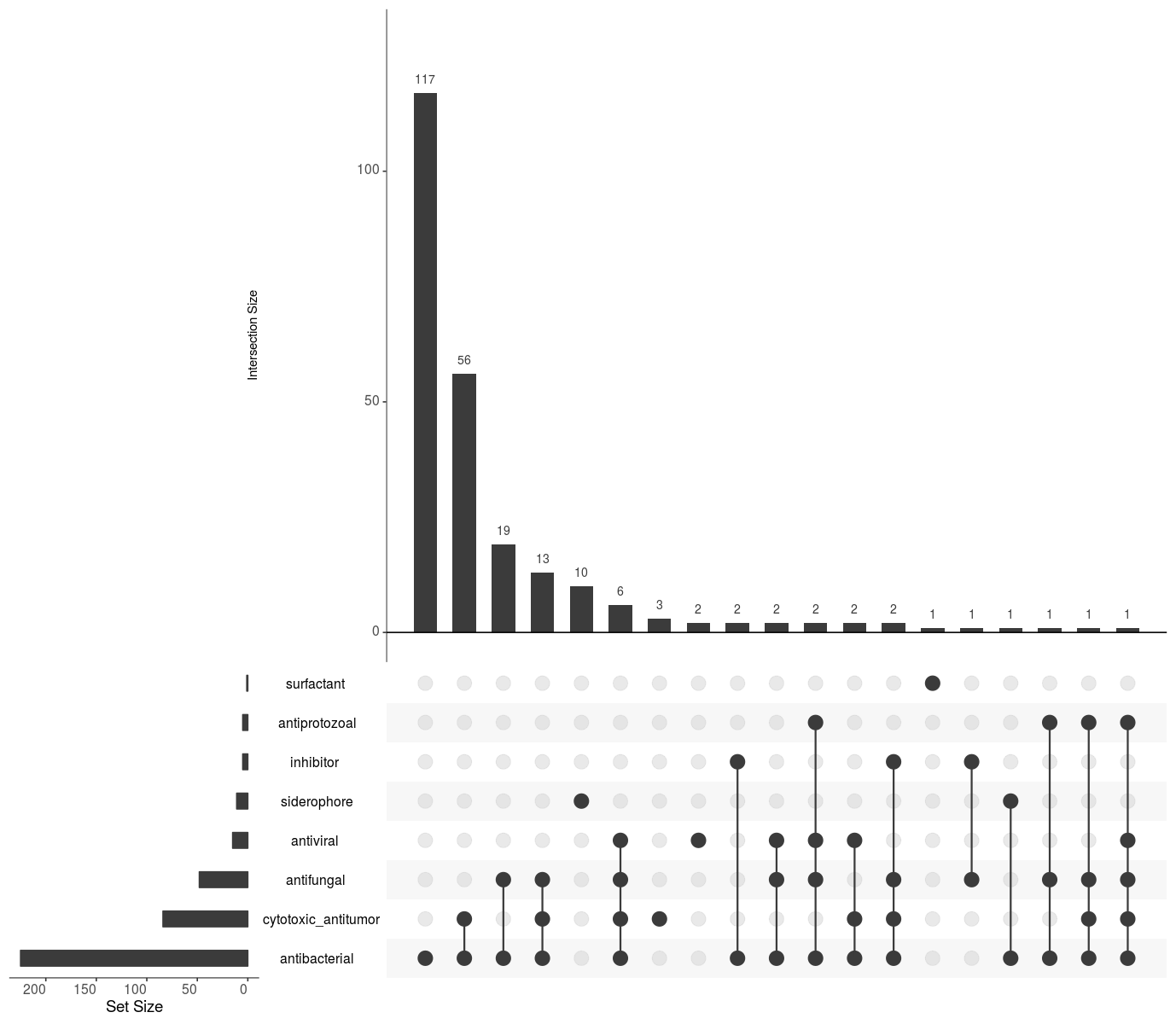

Figure S9: Upset plot showing intersections of NPBdetect predicted bioactivities among BGCs. Vertical bars represent shared bioactivity combinations, while horizontal bars indicate the total number of BGCs per bioactivity class.

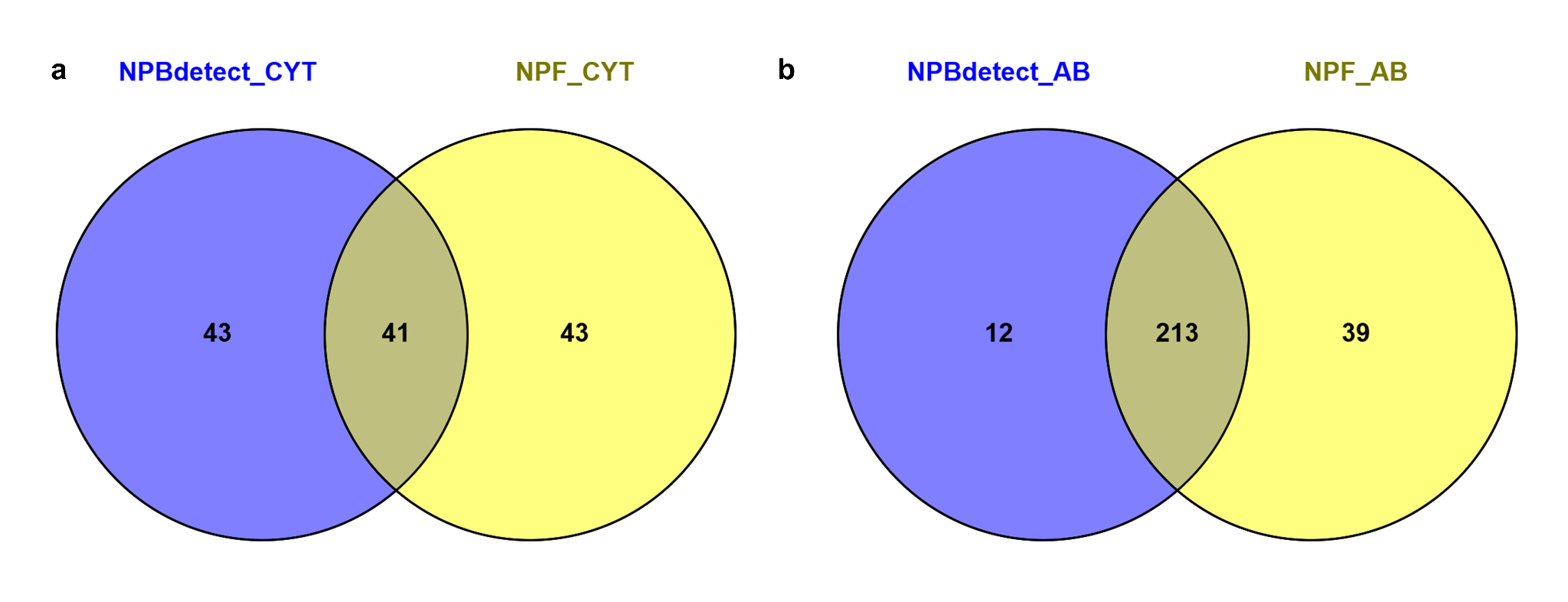

Figure S10: Venn diagrams showing overlap of BGCs predicted to be a) antibacterial (AB) and b) cytotoxic/antitumor (CYT) by two bioactivity prediction tools, NPF and NPBdetect, strengthening the confidence in predicted bioactivities.

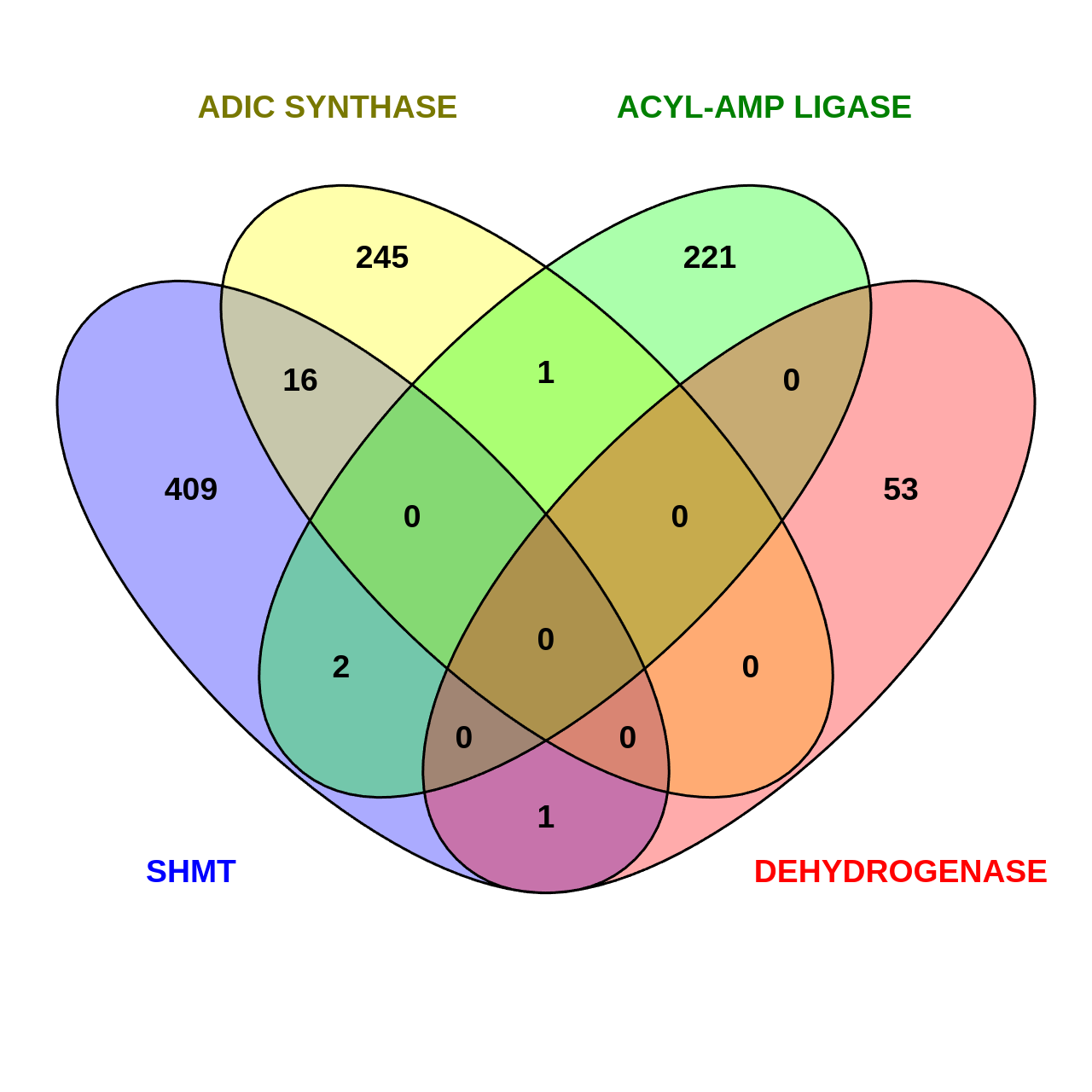

Figure S11: Screening ABC-HuMi database for putative benzoxazolinate-like BGCs: overlap analysis of core biosynthetic protein homologs of benzoxazolinate-like BGCs identified in MAGs sourced from the ABC-HuMi database revealed no overlapping metagenomic regions containing the required combination of protein homologs, indicating the absence of clusters in the analysed MAGs.

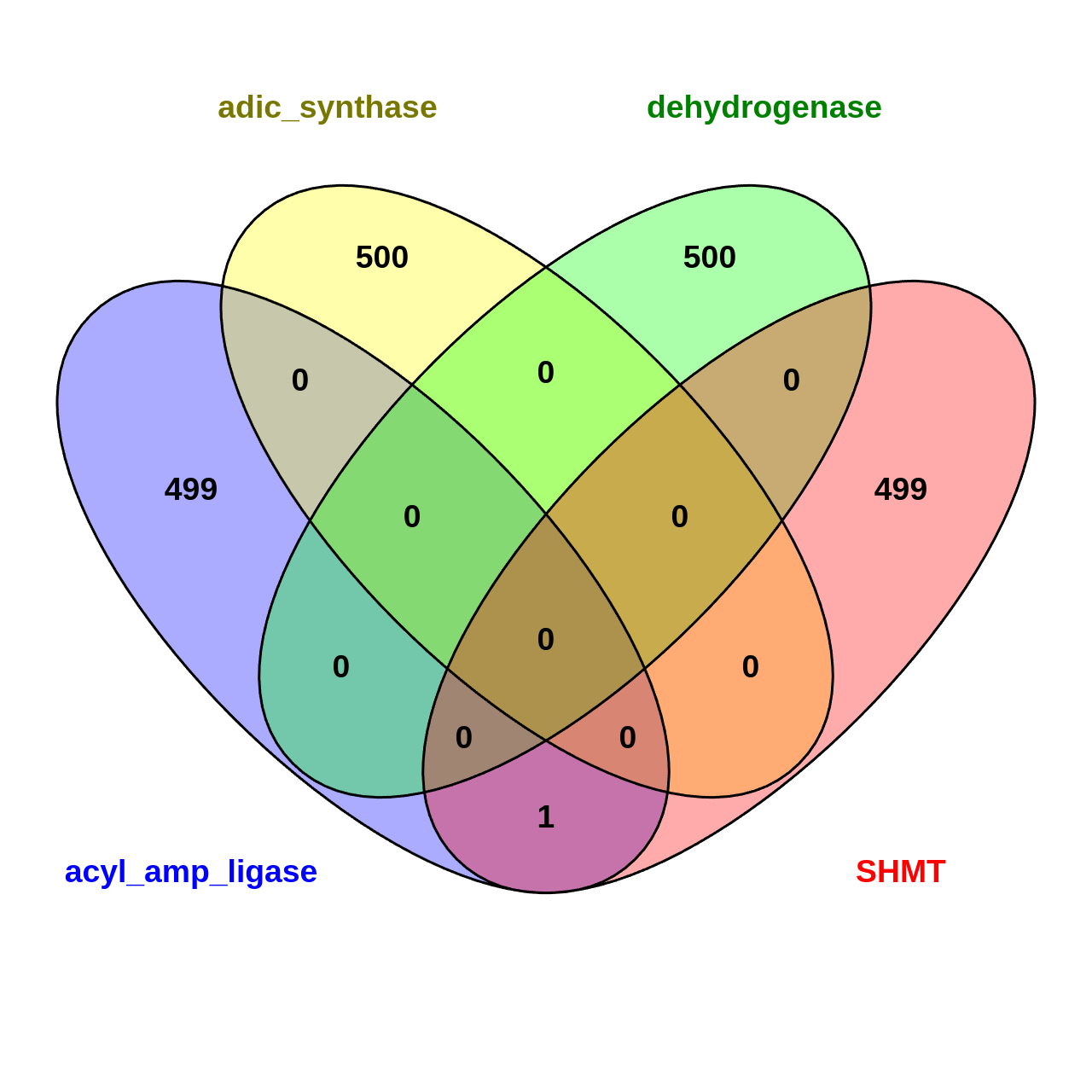

Figure S12: Screening BGC atlas database for putative benzoxazolinate-like BGCs: overlap analysis of core biosynthetic protein homologs of benzoxazolinate-like BGCs identified in MAGs sourced from the BGC Atlas database revealed no overlapping metagenomic regions containing the required combination of protein homologs, indicating the absence of clusters in the analysed MAGs.

Figure S13: Profile HMMs for proposed antiSMASH detection rules for benzoxazolinate-associated clusters

**Conditions for benzoxazolinate-like**

NRPS-like: cds((PP-binding or NAD_binding_4) and (AMP-binding or A-OX) and not alpha_am_amid)

NRPS: cds(Condensation and (AMP-binding or A-OX) and not alpha_am_amid)

**Core**

cds ((FMN_red), (GATase and chorismate_bind))

**Conditions for benzobactin -like**

NRPS: cds(Condensation and (AMP-binding or A-OX) and not alpha_am_amid),

**Core**:

((FMN_red), and (GATase and chorismate_bind) and (SHMT), and (MFS_1))

**Conditions for ashimide-like**

NRPS: cds(Condensation and (AMP-binding or A-OX) and not alpha_am_amid)

NRPS-like: cds((PP-binding or NAD_binding_4) and (AMP-binding or A-OX) and not alpha_am_amid)

**Core**:

cds ((FMN_red) and (GATase, and chorismate_bind) and (MFS_1) and (SHMT) and (p450) and (Methyltransf_11 or Methyltransf_19))

**In some benzoxazolinate-associated BGCs, the ADIC synthase (*phzE*) is harbored from the phenazine biosynthesis cluster.**

**New distance rule:**

Identify the *phzE* at a distance of ≥ 2 Megabase (Mb)
